# Multiple Ciliary Localization Signals Control INPP5E Ciliary Targeting

**DOI:** 10.1101/2022.01.25.477585

**Authors:** Dario Cilleros-Rodriguez, Raquel Martin-Morales, Pablo Barbeito, Abhijit Deb Roy, Abdelhalim Loukil, Belen Sierra-Rodero, Gonzalo Herranz, Olatz Pampliega, Modesto Redrejo-Rodriguez, Sarah C. Goetz, Manuel Izquierdo, Takanari Inoue, Francesc R. Garcia-Gonzalo

## Abstract

Primary cilia are sensory membrane protrusions whose dysfunction causes diseases named ciliopathies. INPP5E is a ciliary phosphoinositide phosphatase mutated in ciliopathies like Joubert syndrome. INPP5E regulates numerous ciliary functions, such as cilium stability, trafficking, signaling, or exovesicle release. Despite its key ciliary roles, how INPP5E accumulates in cilia remains poorly understood. Herein, we show that INPP5E ciliary targeting requires its folded catalytic domain and is controlled by four ciliary localization signals (CLSs), the first two of which we newly discover: LLxPIR motif (CLS1), W383 (CLS2), FDRxLYL motif (CLS3) and CaaX box (CLS4). We answer two long-standing questions in the field. First, partial redundancy between CLS1 and CLS4 explains why CLS4 is dispensable for ciliary targeting. Second, the essential need for CLS2 on the catalytic domain surface clarifies why CLS3 and CLS4 are together insufficient for ciliary accumulation. Furthermore, we reveal that some Joubert syndrome mutations in INPP5E catalytic domain affect its ciliary targeting, and shed light on the mechanisms of action of each CLS. Thus, we find that CLS2 and CLS3 promote interaction with TULP3 and ARL13B, while downregulating CEP164 binding. On the other hand, CLS4 recruits PDE6D, RPGR and ARL13B, and cooperates with CLS1 in ATG16L1 binding. Lastly, we show INPP5E immune synapse targeting is CLS-independent. Altogether, we reveal unusual complexity in INPP5E ciliary targeting mechanisms, likely reflecting its multiple key roles in ciliary biology.

## INTRODUCTION

Primary cilia are solitary membrane protrusions acting as cellular antennae. They emanate from the basal body, a specialized mother centriole, and consist of a microtubule shaft, or axoneme, surrounded by the ciliary membrane, which is topologically continuous with, but compositionally distinct from, the plasma membrane (PM). For cilia to perform their signaling functions, they must accumulate specific receptors and signal transducers. For this to happen, these proteins must first reach the ciliary base, from where they can enter cilia by crossing the transition zone (TZ), the border region separating the ciliary compartment from the rest of the cell. If they can make it inside cilia, the ciliary levels of these proteins will then depend on the balance between ciliary entry and exit rates, a balance that can shift over time. Ciliary entry and exit rates in turn depend on how proteins interact with TZ components, and on whether they associate with specialized ciliary trafficking machinery, such as intraflagellar transport (IFT) trains, microtubule motor-driven multiprotein assemblies whose components, like the IFT-B, IFT-A and BBSome complexes, selectively bind ciliary cargoes to mediate their transport into or out of cilia ^1–3^.

Ciliary malfunction causes ciliopathies, a diverse group of human diseases, many of which are rare autosomal recessive syndromes. One such disease is Joubert syndrome (JBTS), affecting ≈ 1 in 100,000 people worldwide and whose pathognomonic signature is the molar tooth sign (MTS), a cerebellar and midbrain malformation observable by magnetic resonance imaging (MRI). JBTS patients may also present with mild to severe intellectual disability, hypotonia, ataxia, oculomotor apraxia, apnea/hyperpnea, polydactyly, kidney cysts and retinal dystrophy. Genetically, JBTS can be caused by mutations in over 30 different genes, all involved in ciliary function ^2,4–6^.

More specifically, JBTS-causative genes regulate a ciliary signaling network, one of whose main nodes is the ciliary phosphoinositide phosphatase INPP5E (Inositol polyphosphate-5-phosphatase E, formerly known as Pharbin, or as type IV 72 kDa 5-phosphatase) ^7–11^. Evidence for the central role of INPP5E in JBTS includes: (i) INPP5E is one of the most commonly mutated JBTS genes ^6^; (ii) mouse models of INPP5E loss of function recapitulate key features of human JBTS, including the axon tract defects leading to MTS in humans ^7,12^; (iii) most other JBTS genes encode proteins required for INPP5E ciliary targeting ^3,5,13–20^; and (iv) the few JBTS genes that may not be needed for INPP5E ciliary targeting regulate the same pathways as INPP5E, like Hedgehog (Hh) signaling or ciliary stability ^21–27^.

Almost all JBTS-causing INPP5E mutations are missense mutations affecting its catalytic domain ^6,28,29^. In all cases tested, these mutations impaired 5-phosphatase activity towards one or both of its main substrates, PI(4,5)P2 and PIP3 8,29. INPP5E mutations can also cause other ciliopathies, including retinitis pigmentosa (RP), Leber congenital amaurosis (LCA), and MORM syndrome (Mental retardation, Obesity, Retinal dystrophy and Micropenis in males). Unlike JBTS, RP and LCA, the MORM mutation does not affect the catalytic domain, instead removing INPP5E’s C-terminal CaaX box, whose farnesylation tethers INPP5E to membrane ^7,29–31^. Besides its catalytic domain and C-terminus, INPP5E also contains a proline-rich N-terminal region, where no ciliopathy mutations have been reported ^7–11,29^.

INPP5E plays multiple important roles at the cilium. Among others, these roles include regulation of: (i) ciliary phosphoinositide levels, (ii) ciliary protein composition, (iii) ciliary Hedgehog and PI3K signaling, (iv) ciliary ectovesicle release, (v) ciliary stability, and (vi) ciliogenesis ^7,8,12,32–45^. Although most of its functions are ciliary, INPP5E also plays extraciliary roles, like promoting autophagosome-lysosome fusion during autophagy ^46^.

Two motifs in INPP5E protein sequence are known to affect its ciliary targeting: the FDRxLYL motif (aa 609-615) and the CaaX box (aa 641-644), both located in the C-terminal region after the phosphatase domain (aa 297-599) ^7,17,18,47–49^. The CaaX box is not essential for INPP5E ciliary localization, but CaaX box mutants show reduced ciliary targeting, while increasing its proportion at the ciliary base and elsewhere ^7,17,47^. CaaX box farnesylation allows INPP5E to bind phosphodiesterase 6 subunit delta (PDE6D), a prenyl-binding protein that extracts INPP5E from the ciliary base membrane and ferries it across the transition zone. Once inside the cilium, the active form of the monomeric G-protein ADP ribosylation factor (ARF)-like 3 (ARL3) induces dissociation of the PDE6D-INPP5E complex, thus releasing the farnesyl group for insertion into the ciliary membrane. For this to happen, ARL3 must first be activated by a guanine nucleotide exchange factor (GEF) complex consisting of ARF-like 13B (ARL13B), an atypical small G-protein, and its cofactor Binder of ARL2 (BART) ^17,18,48,50–53^.

Intriguingly, despite INPP5E farnesylation not being essential for ciliary targeting, the latter is completely dependent on PDE6D, ARL3 and ARL13B, all of them also JBTS-causative genes ^5,16–18^. Although the reason for this apparent discrepancy is unknown, part of the answer might relate to the retinitis pigmentosa GTPase regulator (RPGR), an RP-associated protein required for INPP5E ciliary targeting, and whose own ciliary localization depends on its geranylgeranylated CaaX box binding to PDE6D ^54–60^.

Regarding ARL13B, its direct interaction with INPP5E probably explains its strong requirement for INPP5E ciliary targeting ^18,48,49^. This interaction is mediated by the FDRxLYL motif, which, unlike the CaaX box, is absolutely required for INPP5E ciliary localization ^18^. Ciliary localization of ARL13B is in turn dependent on Tubby-like protein 3 (TULP3), a phosphoinositide-binding adaptor that links ciliary membrane cargoes to IFT trains ^34,61–64^. This probably explains why TULP3 is also required for INPP5E ciliary targeting, although a more direct connection between TULP3 and INPP5E might also exist ^61,64^. Other proteins needed for INPP5E ciliary targeting include the centrosomal protein of 164 kDa (CEP164), which is involved in INPP5E recruitment to the ciliary base ^18,39,65–67^, and the autophagy-related protein 16-like 1 (ATG16L1), also implicated in ciliogenesis and ciliary trafficking^68,69^.

Although the C-terminal region of INPP5E contains both FDRxLYL motif and CaaX box, this region alone is not sufficient to target INPP5E to cilia. In contrast, a fragment containing both phosphatase domain and C-terminal region suffices for ciliary accumulation ^18^. This indicates that something in or near the catalytic domain is also essential for ciliary targeting, but the reasons for this requirement are unknown.

Herein, we start by elucidating why the catalytic domain is required for ciliary targeting. There are two reasons for this: (i) the FDRxLYL motif is part of the catalytic domain’s globular fold, even though the motif is outside the conserved domain as defined by primary sequence analysis; and (ii) a key catalytic domain residue, W383, which is physically adjacent to the FDRxLYL motif in the domain’s crystal structure, is also specifically and strongly required for ciliary accumulation. We then resolve another lingering question in the field: why the CaaX box is dispensable for INPP5E ciliary localization. We show that the CaaX box is partially redundant with the LLxPIR motif, located near the end of the N-terminal region. Thus, while single deletion of CaaX box or LLxPIR moderately reduces ciliary targeting, simultaneous deletion of both completely abolishes it. Therefore, we reveal that INPP5E ciliary accumulation depends on the interplay between four different ciliary localization signals (CLS1-4), the first two of which we newly identify here: the LLxPIR motif (CLS1), W383 (CLS2), the FDRxLYL motif (CLS3) and the CaaX box (CLS4). In the second half of this work, we systematically examine how each of these CLSs affects INPP5E interactions with proteins required for its ciliary targeting. Through this approach, we find that CLS2 and CLS3 function by promoting interaction with TULP3 and ARL13B, while also antagonizing CEP164 binding. On the other hand, CLS4 is needed for association to PDE6D, RPGR and, in cooperation with CLS1, to ATG16L1. Altogether, our data represent a major step forward in the field and reveal an unprecedented degree of complexity in the ciliary targeting mechanisms of INPP5E, as compared to other known ciliary cargoes, for which a single or at most two CLSs suffice to explain ciliary accumulation ^1,70–73^.

## RESULTS

### INPP5E catalytic domain encompasses the FDRxLYL motif and is required for ciliary targeting

Ciliary accumulation of human INPP5E requires the conserved FDRxLYL motif (aa 609-615) ^18^. This motif lies downstream of INPP5E’s highly conserved phosphatase domain, as defined by the InterPro protein signature database (InterPro domain IPR000300, aa 297-599) ^18,74^. However, INPP5E crystal structure reveals a more extensive globular phosphatase domain, spanning residues 282-623, on whose surface the FDRxLYL motif folds (PDB ID: 2xsw) (**Fig.1a**) ^75^. Indeed, the crystallographic data show that the FDRxLYL motif, and the alpha-helix it is nested in, interact with several other catalytic domain residues (such interactions include D610-L362, Y614-P358, R620-E347 and R621-E354). Recently, a 3D model of full length INPP5E was generated using AlphaFold, a remarkably accurate machine learning algorithm for protein structure prediction ^76^. The AlphaFold model closely matches the crystal structure, including the FDRxLYL motif’s structure and location. Thus, based on the structural data, we conclude that INPP5E’s catalytic domain spans residues 282-623, and therefore encompasses the FDRxLYL motif, which lies on the domain’s surface. Accordingly, from now on, when we speak of the catalytic or phosphatase domain, we will be referring to the globular domain spanning residues 282-623, unless we specify otherwise.

**Figure 1.**
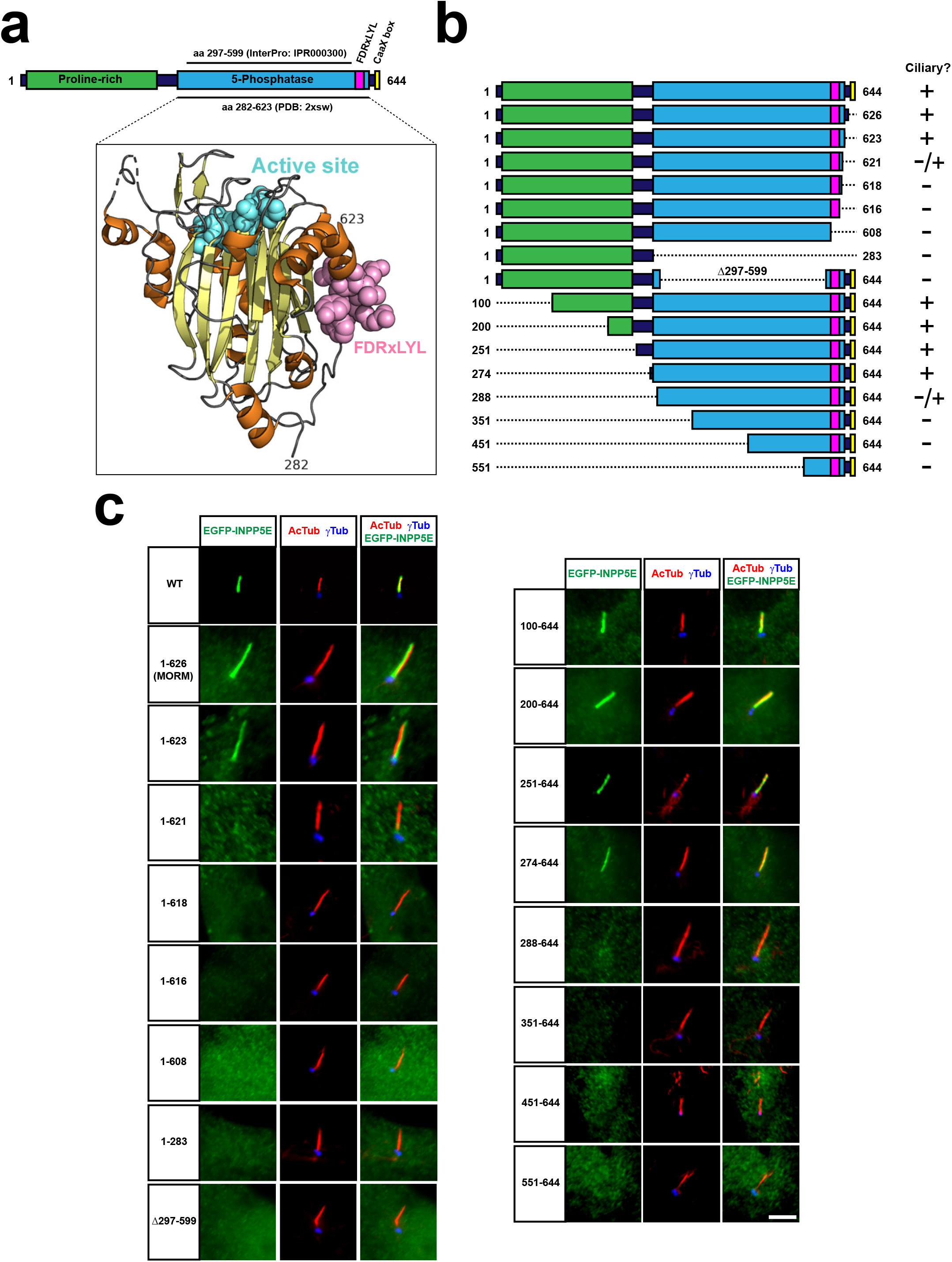
INPP5E catalytic domain encompasses FDRxLYL motif and is required for ciliary targeting. **(a)** Top diagram represents full length human INPP5E protein sequence (aa 1-644). Depicted are the proline-rich region (aa 10-242, Uniprot), the previously reported ciliary localization signal (FDRxLYL, aa 609-615 ^18^), and the CaaX box driving farnesylation (aa 641-644). Also shown is the inositol polyphosphate 5-phosphatase catalytic domain, whose most conserved core corresponds to InterPro domain IPR000300 (aa 297-599), but which actually spans aa 282-623, as revealed by its crystal structure, available at the Protein Data Bank (PDB) and displayed below (PDB ID: 2xsw). Notice how FDRxLYL residues (in magenta above and below) are part of the catalytic domain, on whose surface they fold. The 3D structure also shows active site residues in cyan, alpha-helices in orange, and beta-strands in yellow (including the beta-sandwich at the domain’s core, and a small beta-hairpin near the active site). **(b)** Schematic representation of full length human INPP5E (1-644) and its deletion mutants used in this figure, indicating on the right which ones localize to cilia. **(c)** Representative immunofluorescence images of cilia from hTERT-RPE1 cells transfected with the indicated EGFP-INPP5E constructs. Cells were stained with antibodies against acetylated α-tubulin (AcTub), γ-tubulin (γTub) and EGFP to detect the fusion proteins. Scale bar, 5 μm.

To clarify how INPP5E’s phosphatase domain controls ciliary targeting, we generated a series of INPP5E deletion mutants lacking different portions of the N or C-terminal regions, or lacking only the InterPro-defined catalytic domain (**Fig.1b**). Interestingly, INPP5E ciliary accumulation closely correlated with the presence of an intact structurally-defined catalytic domain (aa 282-623) (**Fig.1b-c**). Deletion of the InterPro-defined catalytic domain (Δ297-599) completely abolished ciliary targeting in EGFP-INPP5E-transfected hTERT-RPE1 cells (**Fig.1c**). N-terminal deletions not affecting the phosphatase domain did not affect ciliary localization, as was the case for constructs 100-644, 200-644, 251-644 and 274-644 (**Fig.1c**). In contrast, 288-644 displayed strongly reduced ciliary targeting, and 351-644, 451-644 and 551-644 completely failed to localize to cilia (**Fig.1c**). C-terminal deletions showed a similar pattern. As previously reported, the MORM mutant (1-626) was found inside cilia and at the ciliary base, and the same was true for 1-623 (**Fig.1c**) ^7^. The 1-621 mutant was only occasionally ciliary, at low levels near the ciliary base (**Fig.1c**). Further deletions from the C-terminus resulted in complete loss of ciliary targeting, as was the case for 1-618, 1-616, 1-608 and 1-283 (**Fig.1c**). Hence, integrity of the structurally-defined catalytic domain is essential for INPP5E ciliary accumulation.

### W383 and FDRxLYL function as specific CLSs on the catalytic domain surface

We first confirmed that, as previously reported, the FDRxLYL motif is essential for INPP5E ciliary targeting (**Fig.2a**) ^18^. Mutation to alanines of both the FDR (aa 609-611) and LYL (aa 613-615) triplets completely abolished ciliary localization (**Fig.2a**). To assess the relative importance of each residue within the FDRxLYL motif, we also made the individual alanine mutants (F609A, D610A, R611A, E612A, L613A, Y614A, L615A). Surprisingly, all of them still localized to cilia, indicating redundancy within the FDR and LYL triplets (**Fig.S1**).

**Figure 2.**
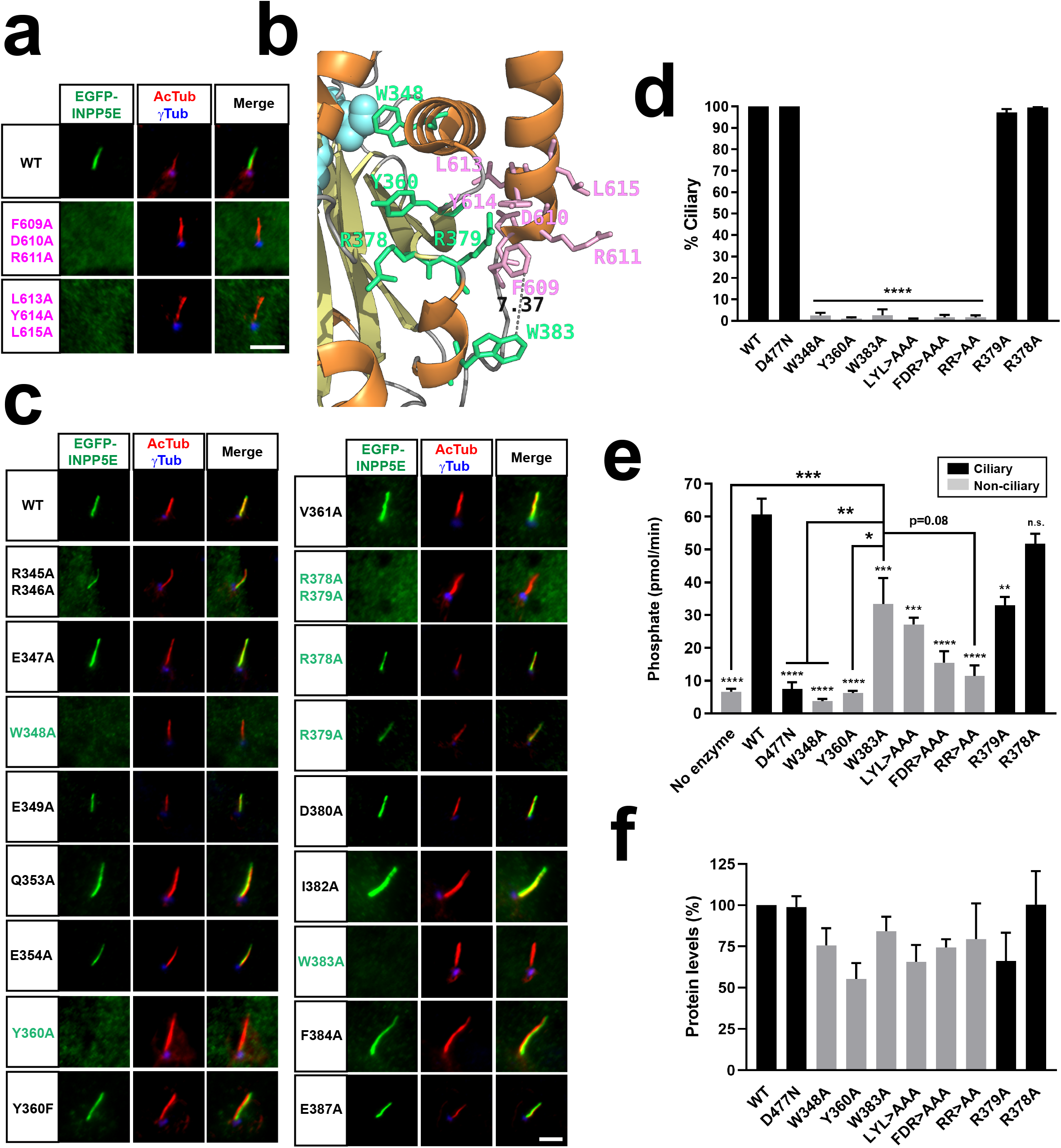
W383 and FDRxLYL motifs act as CLSs on the catalytic domain surface. **(a)** Cilia localization of the indicated FDRxLYL mutants of EGFP-INPP5E was analyzed in hTERT-RPE1 cells as in Figure 1. Scale bars, 5 μm. **(b)** Magnification from INPP5E structure (PDB ID: 2xsw) showing the FDRxLYL motif residues (pink) and adjacent catalytic domain residues shown here to affect ciliary targeting (green). Distance between W383 and F609 is indicated in angstroms. Beta-sheets and alpha-helices are shown as yellow and orange ribbons, respectively. Notice active site region on top left (cyan). **(c)** Cilia localization of the indicated EGFP-INPP5E constructs was analyzed as in (a). Scale bars, 5 μm. **(d)** Percentage of positive cilia was quantitated for each of the indicated constructs. Data are mean±SEM of n=3 independent experiments. Data were analyzed by one-way ANOVA followed by Tukey’s multiple comparisons tests. Significance relative to WT is shown as p<0.0001(****). LYL>AAA: L613A+Y614A+L615A; FDR>AAA: F609A+D610A+R611A; RR>AA: R378A+R379A. **(e)** 5-phosphatase activity, expressed as picomoles of released inorganic phosphate per minute, was measured, using PI(4,5)P2 as substrate, in immunoprecipitates of HEK293T cells transfected with the indicated EGFP-INPP5E variants. Cilia-localized constructs shown as black columns, non-ciliary as grey. Data are mean±SEM of n=9,9,5,4,3,5,3,3,2,3,3 independent experiments (from left to right). Data were analyzed by one-way ANOVA followed by Tukey’s multiple comparisons tests. Significance relative to WT is shown as small asterisks directly above each bar. Significance relative to W383A is shown as bigger asterisks as indicated. In all cases, significance is represented as: p<0.05(*), p<0.01(**), p<0.001(***), p<0.0001(****), or n.s. (not significant). **(f)** Protein levels in the immunoprecipitates used for the activity assays in (e). Western blot bands were quantitated and plotted as percentage of WT. Data are mean±SEM of n=8,5,3,3,5,2,3,3,2,3 independent experiments (from left to right). One-way ANOVA revealed no significant differences.

Given that the FDRxLYL motif lies on the catalytic domain surface (**Fig.1a**), and that catalytic domain integrity is essential for ciliary targeting (**Fig.1b-c**), we hypothesized that INPP5E ciliary targeting relies on a catalytic domain surface including not only the FDRxLYL residues, but also other residues whose proximity to FDRxLYL is dependent upon a folded domain. To test this, we examined the 3D structure of INPP5E’s catalytic domain (PDB ID: 2XSW) in order to identify candidate residues for this putative surface. We did this by selecting residues meeting all or most of the following criteria: (i) located near FDRxLYL motif, on same side of domain; (ii) high exposure to solvent; and (iii) highly conserved in vertebrate INPP5E orthologs. Out of this analysis, we selected sixteen candidate residues: R345, R346, E347, W348, E349, Q353, E354, Y360, V361, R378, R379, D380, I382, W383, F384, E387. Alanine mutation of most of these residues did not affect ciliary targeting of EGFP-INPP5E in hTERT-RPE1 cells (**Fig.2b-c**). Exceptions were W348A, Y360A, W383A and R378A+R379A, all of which completely failed to accumulate in cilia (**Fig.2c-d**). The R378 and R379 arginines are redundant, as the double but not single mutations impeded cilia localization (**Fig.2c-d**).

Mistargeting of these four mutants could be due to loss of catalytic domain integrity, which is critical for ciliary targeting (**Fig.1**). If so, then phosphatase activity would also be disrupted in these mutants. To test this, we immunoprecipitated these EGFP-INPP5E mutants from transfected HEK293T lysates and measured their PI(4,5)P2 5-phosphatase activity in the immunoprecipitates (IPs). EGFP-INPP5E wild type was used as positive control, whereas negative controls included: (i) a reaction with substrate but no enzyme, to assess the rate of basal PI(4,5)P2 dephosphorylation; and (ii) the catalytically inactive D477N INPP5E mutant, lacking a critical active site aspartate but normally localizing to cilia ^8,11,32,77,78^.

Compared to negative controls, phosphate release by WT was about 12-fold faster, a very significant difference (**Fig.2e**). W348A and Y360A were completely inactive, and R378A+R379A nearly so (**Fig.2e**). Hence, lack of ciliary targeting in these mutants may be non-specific. Unlike R378A+R379A, the cilia-localized R378A and R379A single mutants retained activity, either fully (R378A) or partially (R379A). Like R379A, W383A was half as active as WT, with a very significant ≈6-fold increase over negative controls (**Fig.2e**). The same was true for the LYL triplet mutant, whereas the FDR mutant was somewhat less active, with a ≈3-fold increase over controls (**Fig.2a-e**). Quantitation by anti-EGFP immunoblot of the protein levels of each mutant in the IPs used to measure activities revealed no significant differences for any of the mutants, suggesting that activity differences mostly reflect changes in intrinsic enzyme properties, rather than changes in protein stability (**Fig.2f**).

These data suggest that W383 and FDRxLYL function as *bona fide* CLSs, as their strict requirement for ciliary targeting cannot fully be accounted for by their moderate effects on enzyme activity. Accordingly, R379A, a mutation immediately adjacent to FDRxLYL and W383, displayed the same moderate effects on activity, but had no effect on ciliary localization (**Fig.2b-f**).

Still, the partial activity loss in W383A and FDRxLYL mutants suggests that catalytic domain integrity may be partially lost as well. To examine this further, we assessed the half-lives of these mutants in HEK293T cells using the protein translation inhibitor cycloheximide (**Fig.S2**). After 5, 10 or 24 hours in cycloheximide, EGFP-INPP5E(WT) levels were virtually unaffected, indicating a long half-life of several days. For W383A and the triplet FDR and LYL mutants, initial levels were not significantly different from WT (**Fig.S2a-b**). From those initial levels (100%), W383A fell to 60% in the first 10 hours, but then remained at 60% by 24 hours, rather than diminishing further (**Fig.S2c**). Similar results were seen for the FDR mutant, whereas LYL’s curve ran closer to WT’s (**Fig.S2c**).

Thus, W383A showed a biphasic kinetics, whose most parsimonious explanation appears to be the existence of two protein populations: ≈40% of W383A would be unstable with a half-life of ≈5 hours, whereas the remaining ≈60% would be stable, with a half-life of days, like WT (**Fig.S2c**). Presumably, the unstable form would not be properly folded and would be inactive, whereas the stable form would be folded and active, which would explain why W383A reduces activity ≈2-fold (**Fig.2e**). Although this model remains speculative, the fact that ≈50-60% of W383A is stable and enzymatically active supports the idea that W383 is a *bona fide* CLS, since loss of ciliary targeting in W383A is virtually complete, and much stronger than a 2-fold reduction (**Fig.2c-d**). Similar points support the specificity of FDRxLYL as CLS.

We also explored how W383 substitution to residues other than alanine affects ciliary targeting and enzyme activity (**Fig.S3**). To do this, we mutated W383 to another aromatic residue (W383F), to several aliphatic residues (W383I, W383L, W383M, W383V), or to acid (W383E) or basic (W383R) residues. Interestingly, W383F was fully active and ciliary, indicating that an aromatic ring at this position suffices for both functions (**Fig.S3**). In contrast, all other mutations fully suppressed ciliary targeting, like W383A (**Fig.S3a**). Interestingly, enzyme activity of W383I, W383L, W383M and W383V was the same as for W383A, suggesting that an aromatic ring in this position is important for both targeting and activity (**Fig.S3b-c**). Overall, the data in this section show that W383 and FDRxLYL, despite their moderate effects on enzyme activity, function as specific CLSs to target INPP5E to cilia.

### The LLxPIR motif cooperates with the CaaX box to target INPP5E to cilia

Besides W383 and FDRxLYL, the C-terminal CaaX box (641-CSVS-644) also modulates INPP5E ciliary targeting. Although CaaX box deletion causes mistargeting of some INPP5E molecules to the ciliary base or other cellular membranes, CaaX box mutants still accumulate in cilia (**Fig.1c**) ^7,17,47^. This is somewhat puzzling, as INPP5E ciliary targeting strongly depends on the farnesyl receptor PDE6D ^17^.

In the course of our experiments, we made a serendipitous observation that led us to novel insights into how non-farnesylated INPP5E manages to still accumulate in cilia. Initially, we found that INPP5E’s N-terminus (aa 1-273), while being dispensable for ciliary targeting of full length INPP5E (**Fig.1b-c**), is required for ciliary targeting of INPP5E’s MORM mutant (Δ627-644) (**Fig.3e**). We subsequently saw that this effect of the N-terminus is mediated by residues 251-273, and that these residues are strongly required for ciliary targeting of the C641S mutant, in which the farnesylated CaaX box cysteine is replaced by non-farnesylatable serine (**Fig.3a-c**). Thus, while both Δ251-273 and C641S single mutants localized inside cilia, the double (Δ251-273)+C641S mutant completely failed to do so, despite accumulating at the ciliary base (**Fig.3a-c**). Interestingly, upon careful observation and quantitation, both single mutants already showed a partial loss of intraciliary targeting, aside from their ciliary base accumulation (**Fig.3a-c**). Thus, the single Δ251-273 mutant behaves as previously reported for CaaX mutants (and as shown here for C641S), except that Δ251-273 reduces ciliary targeting even more than C641S (**Fig.3c**). These data indicate that the CaaX box and residues 251-273 cooperate to target INPP5E to cilia: not only are both sequences required for optimal ciliary targeting of the wild type protein, but their functions are partially redundant, each becoming essential when the other one is missing (**Fig.3a-c**).

**Figure 3.**
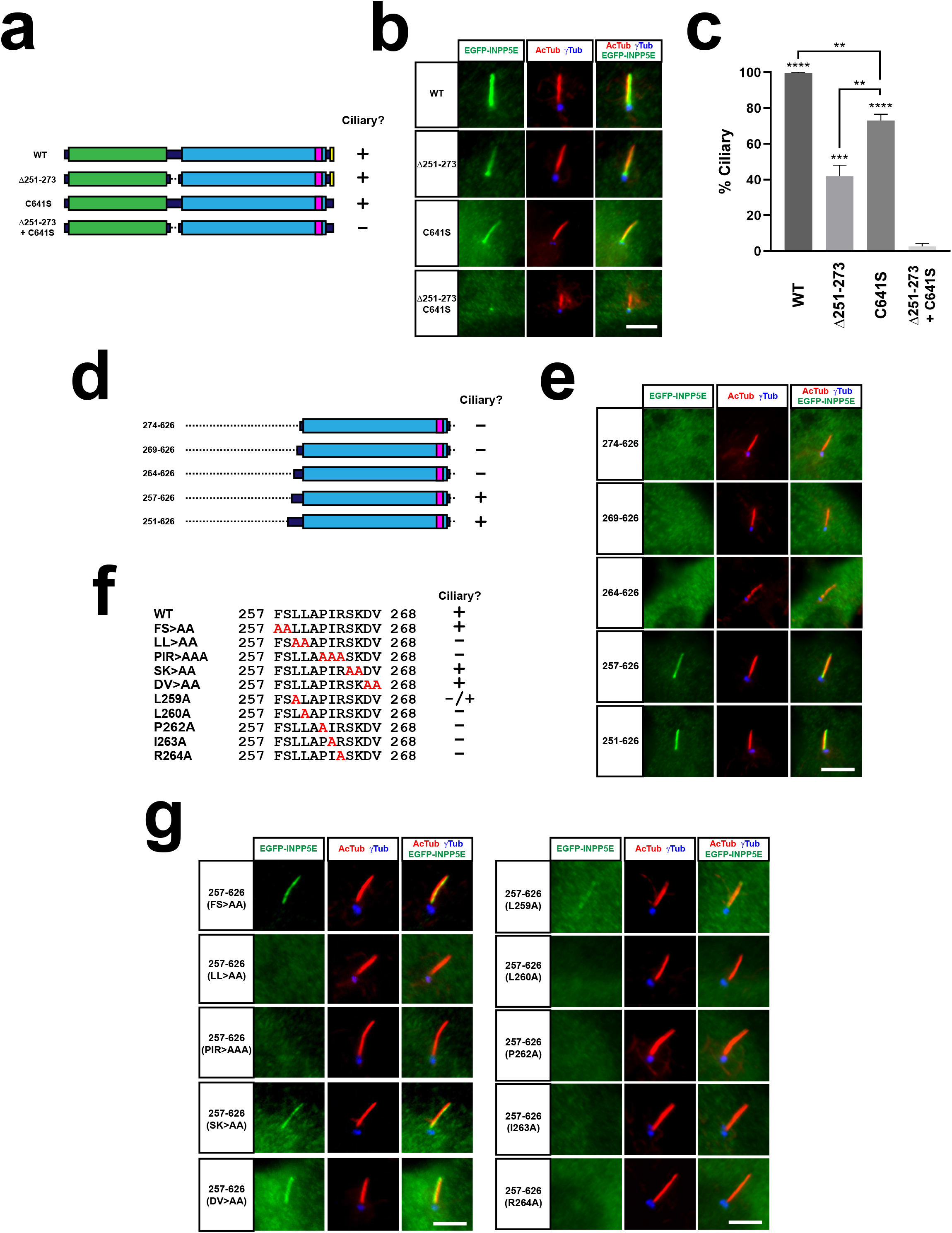
The LLxPIR motif is a novel CLS that cooperates with the CaaX box to mediate INPP5E ciliary targeting. **(a)** Schema of full length human INPP5E and its mutants used in (b-c). Cilia localization of each mutant is indicated on the right. **(b)** Cilia localization of WT and indicated mutants was analyzed in hTERT-RPE1 cells as in Figures 1-2. Scale bar, 5μm. **(c)** Quantitation of data from (b). The percentage of positive cilia in transfected cells is shown for the indicated EGFP-INPP5E constructs as mean±SEM of n=3 independent experiments. Data were analyzed by one-way ANOVA with post-hoc Tukey multiple comparisons tests. Statistical significance is depicted as p<0.01(**), p<0.001(***) or p<0.0001(****). Significance is shown relative to Δ251-273+C641S unless otherwise indicated. **(d)** Schema of INPP5E deletion mutants used to map the CLS within aa 251-273. None of these mutants contains the CaaX box (aa 641-644), so their ciliary targeting is strictly dependent on residues 251-273. Cilia localization of each mutant is indicated on the right. **(e)** Cilia localization of the mutants from (d) was analyzed in hTERT-RPE1 cells as in Figures 1-3. **(f)** Sequence of aa 257-268 in wild type INPP5E and indicated mutants, whose ciliary localization in shown on the right. **(g)** Ciliary targeting of INPP5E(257-626) containing the mutations from (f) was analyzed as above. Scale bars, 5μm.

Even though both these mutations are outside the phosphatase domain, it remains possible that their effects on ciliary targeting are due to disruption of catalytic domain integrity. To test this, we measured the PI(4,5)P2 5-phosphatase activity of the aforementioned 274-626 mutant, lacking both the N-terminus (Δ1-273) and the MORM region (Δ627-644). This mutant was fully active, with activity and protein levels indistinguishable from WT (**Fig.S4**). In contrast, the 288-626 mutant, additionally missing residues 275-287 at the beginning of the catalytic domain, had much less activity and reduced levels (**Fig.S4**). Therefore, residues 251-273 and 627-644 are completely dispensable for enzyme activity, indicating that their effects on ciliary targeting are not caused by disruptions in the phosphatase domain. In other words, 251-273 and the CaaX box also behave as *bona fide* CLSs.

We then mapped which residues within 251-273 are responsible for CLS function. To do this, we started with the 274-626 mutant, which fails to accumulate in cilia as mentioned above. To this mutant, we gradually added residues to the N-terminus, thus creating four more mutants: 269-626, 264-626, 257-626 and 251-626 (**Fig.3d**). Of these, 269-626 and 264-626 failed to accumulate in cilia, whereas 257-626 and 251-626 readily did so (**Fig.3e**). Next, starting from 257-626, we generated five alanine-substitution mutants within the 257-FSLLAPIRSKDV-268 region, removing residues FS, LL, PIR, SK and DV, respectively (**Fig.3f**). Of these, residues LL and PIR were essential for ciliary targeting, whereas FS, SK and DV were fully dispensable (**Fig.3f-g**). We then created the single residue mutants spanning the 259-LLAPIR-264 motif. All these mutants (L259A, L260A, P262A, I263A and R264A) abolished cilia localization of 257-626, although weak residual targeting was still observed only for L259A (**Fig.3f-g**). Therefore, the LLxPIR motif is a novel CLS that cooperates with the CaaX box to mediate optimal ciliary targeting of INPP5E.

### INPP5E ciliary targeting is mediated by four conserved CLSs

The above data indicate that INPP5E ciliary targeting depends on four CLSs, which we will heretofore refer to as CLS1 (the LLxPIR motif, aa 259-264), CLS2 (W383), CLS3 (the FDRxLYL motif, aa 609-615), and CLS4 (the CaaX box, aa 641-644). Our data actually show that CLS3 goes beyond the FDRxLYL motif, as deletion of residues 616-621 also interferes with ciliary targeting (**Fig.1b-c**). This is consistent with the original report on CLS3, where residues 619-621 were also shown to modulate ciliary targeting ^18^. Thus, CLS3 spans residues 609-621 in human INPP5E.

Since INPP5E ciliary targeting has been reported in different vertebrate species, we examined whether CLS1-4 are conserved in vertebrate evolution. To do this, we aligned the human INPP5E sequence with those of another mammal (mouse), a bird (crow), a reptile (python), an amphibian (toad) and a fish (zebrafish) (**Fig.4a**). From this analysis, it is clear that all four CLSs are highly conserved in vertebrates. For CLS1, the consensus sequence is [VL]LxPIR, with only the first leucine admitting a conservative change (**Fig.4a**; **Fig.3g**). CLS2’s tryptophan is fully conserved, and so is CLS3, as previously shown (consensus: FDRxLYLxGI[KR]RR) (**Fig.4a**) ^18^. For CLS4 the consensus is C[ST][IV]S, which in all cases encodes a farnesyl transferase-specific CaaX box ^79^. Thus, the four CLSs are highly conserved. Moreover, they are all found on the same side of INPP5E according to the Alphafold model (**Fig.4b**) ^76^.

**Figure 4.**
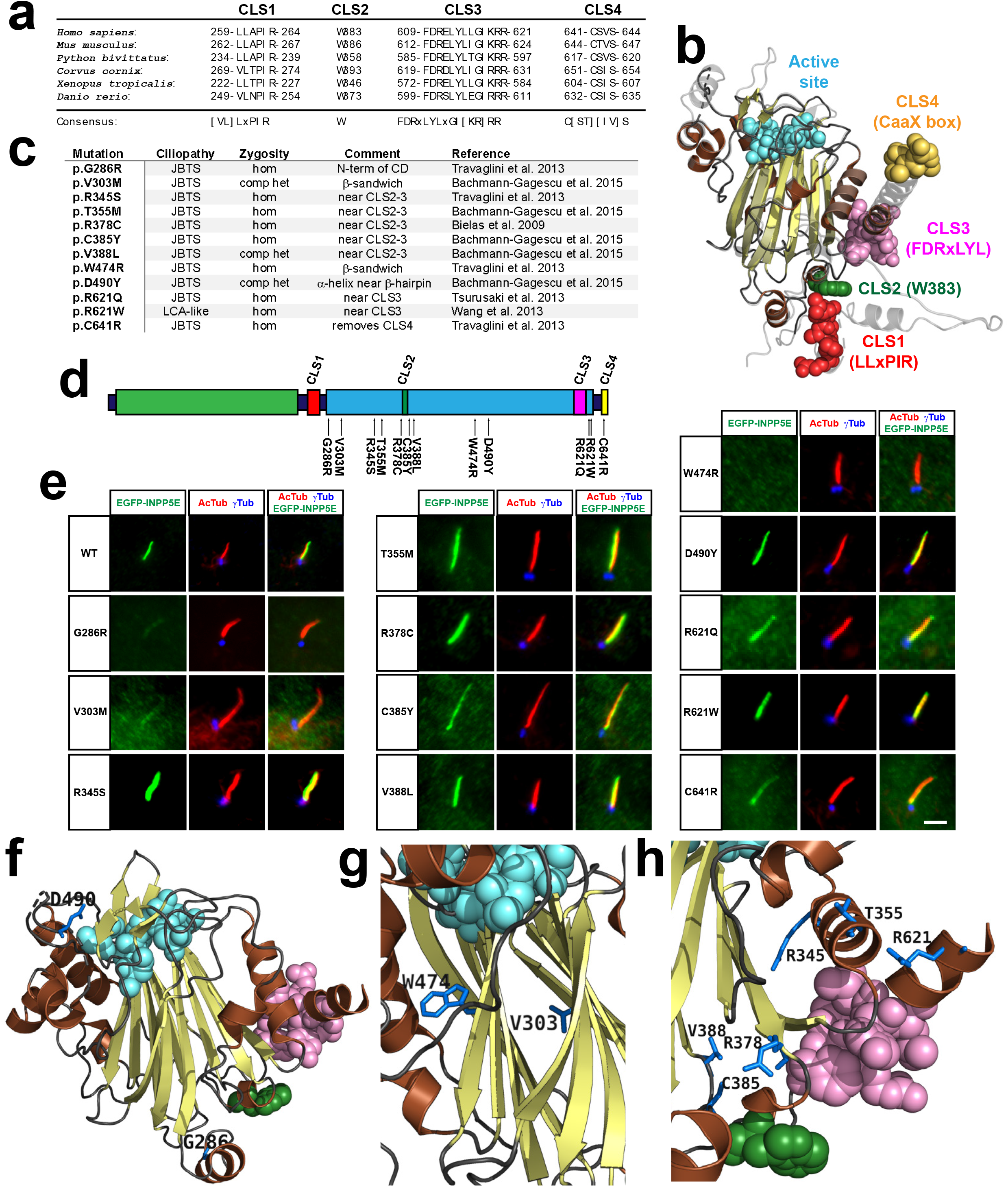
A subset of Joubert syndrome INPP5E mutations abolishes ciliary targeting. **(a)** CLS1-4 are highly evolutionarily conserved in vertebrates, including human (NP_063945.2), mouse (AAH80295.1), python (XP_007441606.1), crow (XP_039417670.1), toad (XP_002935265.1), and zebrafish (NP_001096089.2). Consensus sequences are shown below. **(b)** Alphafold model of INPP5E 3D structure (AF-Q9NRR6-F1) depicting predicted locations of CLS1 (red), CLS2 (green), CLS3 (pink) and CLS4 (yellow). Active site in cyan. Beta-strands and alpha-helices in yellow and brown, respectively. Proline-rich N-terminal region (aa 1-200), predicted to be highly flexible, is not shown. **(c)**Table of INPP5E ciliopathy mutations analyzed here. JBTS: Joubert syndrome; LCA: Leber congenital amaurosis; hom: homozygous; comp het: compound heterozygous. **(d)** Schema of INPP5E protein sequence indicating the locations of the ciliopathy mutations from (c) relative to its four CLSs, its catalytic domain (cyan) and its N-terminal proline-rich region (green). **(e)** Ciliary localization of mutants from (c-d) was analyzed in hTERT-RPE1 cells as in Figures 1-3. Scale bar, 5 μm. (f-h) 3D views of INPP5E catalytic domain (PDB ID: 2xsw) showing the ciliopathy-mutated residues from (c) in dark blue (other colors as in (b)). **(f)** full catalytic domain showing G286 (bottom) and D490 (top left). **(g)** closeup view of beta-sandwich showing W474 and V303. **(h)** closeup view of CLS2-3 region showing R345, T355, R378, C385, V388L and R621.

### Ciliary targeting is affected by some INPP5E ciliopathy mutations

INPP5E ciliary targeting is dependent on CLS1, CLS2, CLS3 and CLS4 (**Figs.2–3**), and on the integrity of its catalytic domain (**Fig.1**). The MORM mutation (Δ627-644), which deletes CLS4 and moderately reduces cilia localization, as mentioned above, is the only known INPP5E ciliopathy mutation affecting its ciliary targeting. However, whether and how other ciliopathy mutations in INPP5E also affect ciliary targeting is largely unknown. To address this, we sought ciliopathy mutations locating near a CLS, or that were likely to compromise catalytic domain integrity. In this manner, we identified twelve mutations, eleven from JBTS and one from LCA-like disease (**Fig.4c-d**) ^6,8,28–31^.

One of the mutations, C641R, replaces the farnesylatable cysteine in CLS4 by an arginine. Not surprisingly, this mutation affected ciliary targeting in the same way as C641S or MORM (**Fig.4d-e**). Several other mutations affected residues located near the CLS2-CLS3 region. This included R345S, T355M, R378C, C385Y, V388L, R621Q and R621W. None of these affected ciliary localization (**Fig.4d-e,h**). Since almost all INPP5E ciliopathy mutations affect the catalytic domain, none were found near CLS1. The closest one to CLS1 was G286R, at the beginning of the catalytic domain. Interestingly, G286R largely abolished ciliary targeting, although this is more likely due to its affecting catalytic domain integrity (**Fig.4d-f**). We also tested mutations, like W474R and V303M, that we thought likely to disrupt the catalytic domain, given their location at the core of the beta-sandwich. Indeed, W474R abolished ciliary targeting, and V303M reduced it considerably (**Fig.4d-e,g**). Since W474R and G286R were found in homozygosis in JBTS patients, this suggests that some JBTS patients cannot target INPP5E to cilia (**Fig.4c,e**). Finally, we also tested the D490Y mutation, located near a beta-hairpin close to the active site, but far from any CLS. D490Y did not affect ciliary targeting, so it probably does not affect domain integrity (**Fig.4d-f**). Altogether, these data show that INPP5E ciliary targeting is sometimes affected in Joubert syndrome, which could contribute to pathogenesis in these cases.

### INPP5E binding to PDE6D is CLS4-dependent

After identifying novel INPP5E CLSs and exploring their role in ciliopathies, we focused on the mechanisms of action of these CLSs. Presumably, these CLSs act by binding to other proteins implicated in INPP5E ciliary targeting. One such protein is PDE6D, a ciliary cargo receptor for prenylated proteins. Although PDE6D binding to INPP5E is CLS4-dependent and CLS3-independent, whether its interaction requires our newly identified CLSs (CLS1 and CLS2) is unknown ^17,18,49,55^. To test this, we cotransfected HEK293T cells with plasmids encoding Flag-PDE6D and EGFP-INPP5E in order to perform coimmunoprecipitation (co-IP) experiments (**Fig.5a**). As expected, Flag-PDE6D robustly co-immunoprecipitated (co-IPed) with EGFP-INPP5E(WT), but not EGFP control (**Fig.5a**). This interaction was completely dependent on CLS4, and completely independent of CLS1, CLS2 and CLS3 (**Fig.5a**). The mutations used to ascertain this were: Δ251-273 (ΔCLS1), W383A (ΔCLS2), F609A+D610A+R611A (ΔCLS3) and C641S (ΔCLS4). The double (Δ251-273)+C641S mutant (ΔCLS1+4) behaved the same as ΔCLS4 (**Fig.5a**). We also tested whether the interaction involved INPP5E’s N-terminal (aa 1-283) or C-terminal (251-644) regions, with the latter being the case for PDE6D (**Fig.5a**).

**Figure 5.**
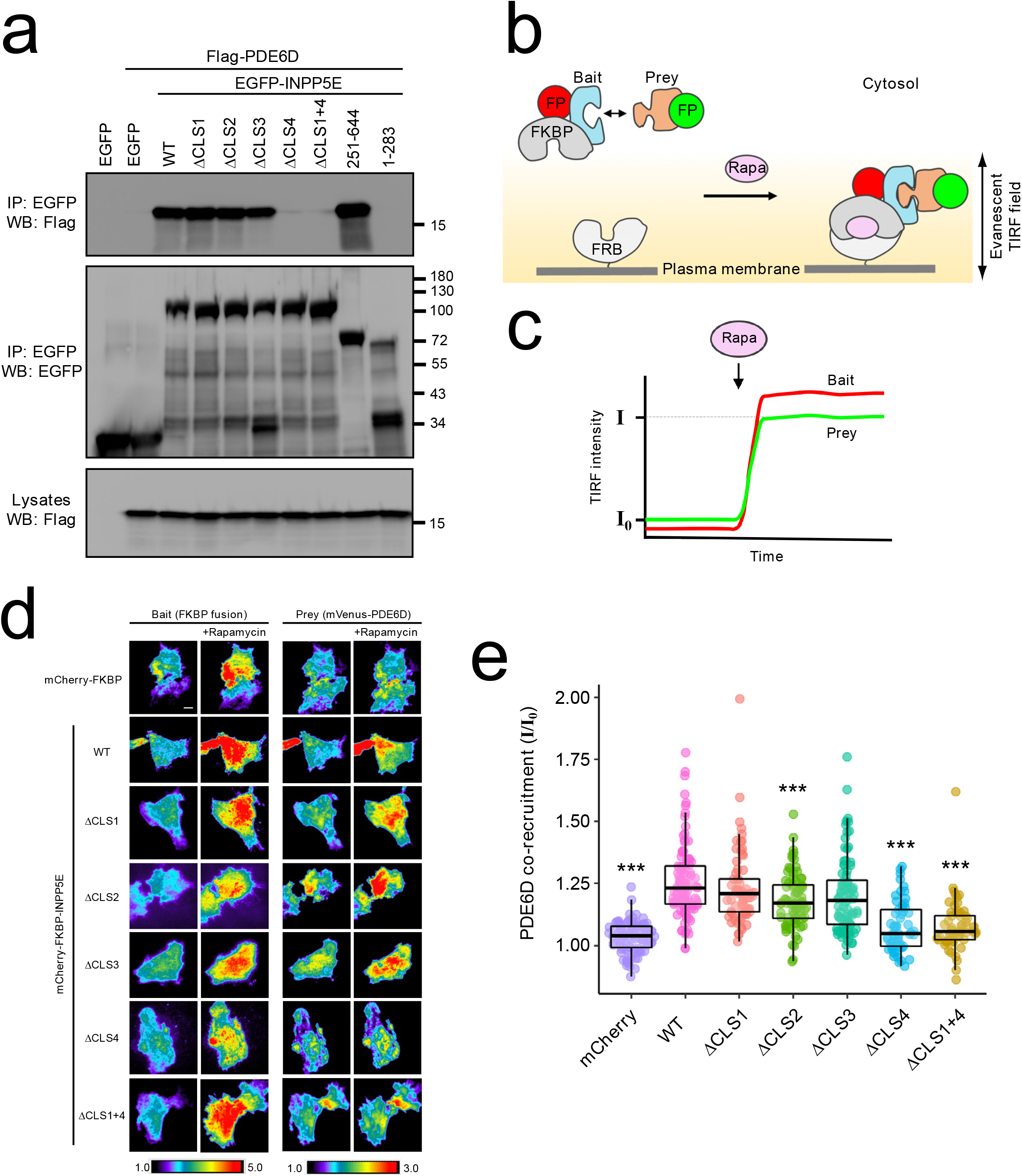
CLS4 promotes INPP5E binding to PDE6D. **(a)** The indicated EGFP-INPP5E variants were coexpressed in HEK293T cells with Flag-PDE6D, as indicated. Lysates were immunoprecipitated with GFP-Trap beads and analyzed by Western blot with the indicated antibodies. **(b)** Schema of chemically-inducible co-recruitment assay. Rapamycin (Rapa)-induced interaction between FKBP and FRB is used to quantitate binding of prey candidates to a bait. FKBP is fused to the prey along with a fluorescent protein, while FRB is tethered to inner leaflet of plasma membrane. Upon rapamycin addition, FKBP binds to FRB, bringing bait (red FP) and associated prey (green FP) to the plasma membrane. **(c)** Recruitment of bait and prey to plasma membrane can be sensitively detected by TIRF microscopy as an increased fluorescence signal. The ratio of final to initial TIRF intensity upon rapamycin addition (I/I0) for the prey provides a quantitative measure of prey’s co-recruitment to plasma membrane by bait, and hence of the prey-bait interaction. **(d)** TIRF microscopy images showing rapamycin-induced plasma membrane recruitment of bait constructs (left) and the corresponding co-recruitment of prey (mVenus-PDE6D, right). Intensity scales are depicted at bottom. Scale bar, 10mm. **(e)** Normalized rapamycin-induced co-recruitment of mVenus-PDE6D (prey) by mCherry-FKBP-INPP5E (WT or indicated mutants), or by mCherry-FKBP (mCherry) as negative control. Individual measurements of n>50 cells per condition are shown. Box and whisker plots represent median, first and third quartiles, and 95% confidence intervals. Statistical significance relative to WT is shown as *** p<0.001 (unpaired Student’s t-tests).

In addition to co-IP, we also studied the INPP5E-PDE6D interaction in vivo by means of co-recruitment assays (**Fig.5b-c**). To do this, we generated bait constructs expressing mCherry-FKBP-INPP5E fusion proteins (WT, ΔCLS1, ΔCLS2, ΔCLS3, ΔCLS4 or ΔCLS1+4), or mCherry-FKBP as negative control. Each of these bait constructs was separately co-expressed in HeLa cells with the prey construct (mVenus-PDE6D), and with FRB-CFP-CaaX, whose prenylated CaaX box tethers FRB to the inner leaflet of the plasma membrane (**Fig.5b**). Upon inducing the FKBP-FRB interaction with rapamycin, this should recruit bait constructs to the plasma membrane, a recruitment that can be monitored and quantitated by total internal reflection fluorescence (TIRF) microscopy ^80,81^. Additionally, if bait and prey interact, then prey co-recruitment to the plasma membrane will also be observed (**Fig.5b-c**). As expected, rapamycin induced robust plasma membrane recruitment of all bait constructs (**Fig.5d**). Also as expected, mVenus-PDE6D co-recruitment was much higher with mCherry-FKBP-INPP5E(WT) than with the mCherry-FKBP control, confirming the specificity of PDE6D-INPP5E binding. As observed in the co-IPs, mVenus-PDE6D co-recruitment was strongly reduced by both the ΔCLS4 and ΔCLS1+4 mutations, indicating a strong CLS4-dependence (**Fig.5d-e**). Also consistent with the co-IPs, ΔCLS1 and ΔCLS3 did not affect the interaction, whereas ΔCLS2 caused a modest reduction in mVenus-PDE6D co-recruitment (**Fig.5d-e**). Altogether, the co-IP and in vivo co-recruitment data demonstrate that CLS4 is the key CLS controlling INPP5E-PDE6D binding.

### INPP5E binding to RPGR is CLS4-dependent

RPGR also interacts with INPP5E and is required for its ciliary targeting ^55^. Moreover, RPGR ciliary targeting is also dependent on PDE6D, which binds to both its geranylgeranylated CaaX box and its RCC1-like domain ^54–60^. Despite all these connections, how RPGR mediates INPP5E ciliary targeting is unclear. To address this, we performed co-IPs between Flag-RPGR and the same EGFP-INPP5E constructs used above for the PDE6D co-IPs. As with PDE6D, the INPP5E-RPGR interaction was abolished by ΔCLS4 and ΔCLS1+4, but was untouched by ΔCLS1, ΔCLS2 or ΔCLS3 (**Fig.S5**). Therefore, INPP5E’s farnesylated CaaX box is also key for its binding to RPGR. Consistently, RPGR strongly interacted with INPP5E’s C-terminal fragment (251-644) but not with the N-terminal one (1-283).

### INPP5E binding to ARL13B is promoted by CLS2, CLS3 and CLS4

ARL13B is another key mediator of INPP5E ciliary targeting ^18,48,49^. ARL13B regulates INPP5E in at least two different ways. First, ARL13B acts as a guanine nucleotide exchange factor (GEF) for ARL3, whose active GTP-bound form promotes dissociation of the PDE6D-INPP5E complex after it reaches the ciliary lumen ^50–53^. Additionally, ARL13B directly interacts with INPP5E and is required for its ciliary retention ^18,48,49^. Since the first mechanism is mediated by ARL3, we checked whether INPP5E binds ARL3. However, we detected no co-IP between EGFP-INPP5E and ARL3-myc, or its constitutively active form ARL3(Q71L)-myc, in accordance with previous data (data not shown) ^18^. Likewise, we found no interaction between INPP5E and BART, a protein that cooperates with ARL13B as a co-GEF for ARL3 (data not shown) ^53^.

We then carried out co-IPs to assess the CLS-dependence of the ARL13B-INPP5E interaction. We readily detected co-IP of endogenous ARL13B with EGFP-INPP5E in HEK293T cells. This co-IP was largely abolished with ΔCLS4 and ΔCLS1+4, and somewhat reduced with ΔCLS2 and ΔCLS3. In contrast, ΔCLS1 had no effect (**Fig.6a**). As with PDE6D and RPGR, INPP5E’s C-terminal region (251-644), but not the N-terminal (1-283), sufficed for the INPP5E-ARL13B interaction (**Fig.6a**). Thus, co-IP experiments indicated that CLS2, CLS3 and CLS4 played a role in ARL13B binding. This was fully corroborated by co-recruitment assays like those in **Fig.5**, in which ARL13B-EYFP was used as prey instead of mVenus-PDE6D (**Fig.6b**). ARL13B-EYFP co-recruitment strongly increased with mCherry-FKBP-INPP5E(WT), as compared to mCherry-FKBP alone, demonstrating a specific interaction. This interaction was unaffected by ΔCLS1 but strongly reduced with ΔCLS2, ΔCLS3, ΔCLS4 and ΔCLS1+4 (**Fig.6b**). Therefore, CLS2, CLS3 and CLS4 all promote INPP5E binding to ARL13B.

**Figure 6.**
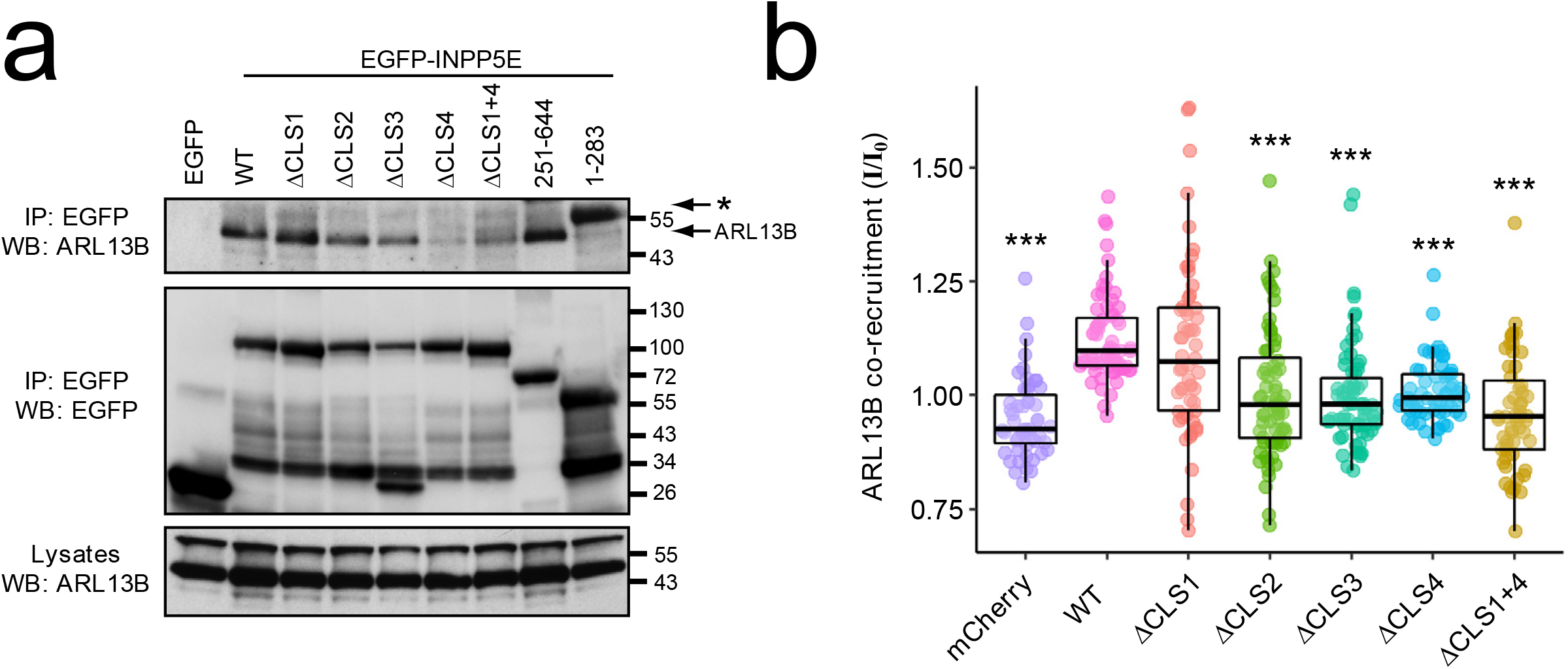
CLS2, CLS3 and CLS4 promote INPP5E binding to ARL13B. **(a)** Coimmunoprecipitation of endogenous ARL13B with the indicated EGFP-INPP5E constructs in HEK293T cells. Asterisk points to EGFP-INPP5E(1-283) band. **(b)** Normalized rapamycin-induced co-recruitment of ARL13B-EYFP (prey) by mCherry-FKBP-INPP5E (WT or indicated mutants), or by mCherry-FKBP (mCherry) as negative control. Individual measurements of n>50 cells per condition are shown. Box and whisker plots represent median, first and third quartiles, and 95% confidence intervals. Statistical significance relative to WT is shown as *** p<0.001 (unpaired Student’s t-tests).

### INPP5E binding to TULP3 is promoted by CLS2 and CLS3

TULP3 is a ciliary trafficking adapter needed for ciliary targeting of membrane proteins such as G protein-coupled receptors, polycystins, ARL13B and INPP5E ^34,61-63,82–84^. However, whether INPP5E and TULP3 interact is not known. We therefore tested this. Indeed, EGFP-INPP5E specifically co-IPed TULP3-myc (**Fig.7a**). Such co-IP was unaffected by ΔCLS1, ΔCLS4 and ΔCLS1+4, but was clearly reduced by ΔCLS2 and ΔCLS3 (**Fig.7a**). Consistently, TULP3 interacted strongly with EGFP-INPP5E(251-644), and much less so with EGFP-INPP5E(1-283) (**Fig.7a**).

**Figure 7.**
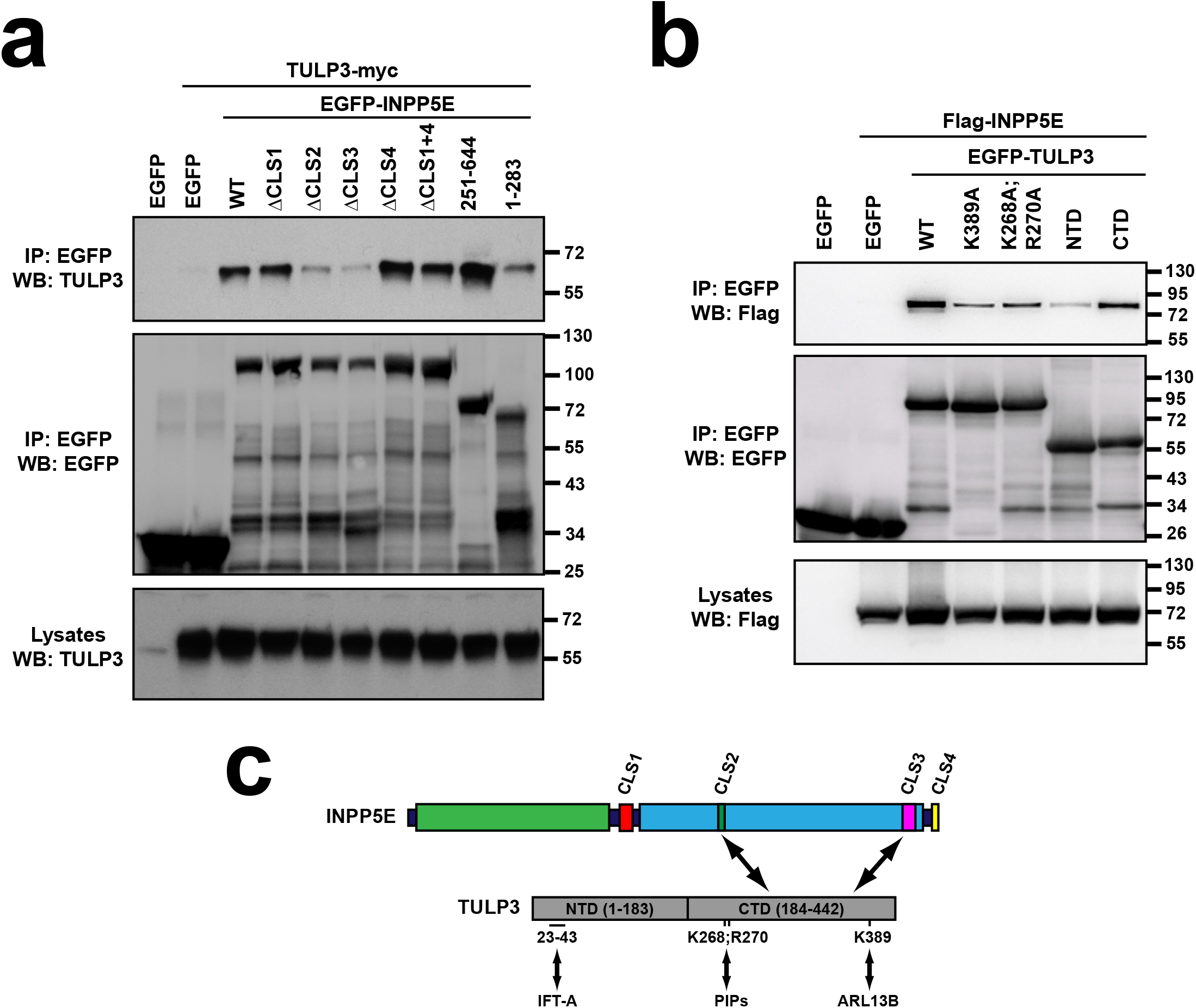
CLS2 and CLS3 promote INPP5E binding to TULP3. **(a)** The indicated EGFP-INPP5E variants were coexpressed in HEK293T cells with TULP3-myc as indicated. Lysates were immunoprecipitated with GFP-Trap beads and analyzed by Western blot with the indicated antibodies. **(b)** The indicated EGFP-TULP3 variants were coexpressed in HEK293T cells with Flag-INPP5E as indicated. Lysates were immunoprecipitated with GFP-Trap beads and analyzed by Western blot with the indicated antibodies. NTD: N-terminal domain (aa 1-183); CTD: C-terminal Tubby domain (aa 184-442). **(c)** Schema of INPP5E-TULP3 interaction. On INPP5E’s side, the interaction mostly involves the catalytic domain, requiring CLS2 and CLS3. On TULP3’s side, the interaction occurs mostly through the CTD and is affected by the ARL13B-binding K389, and by the phosphoinositide (PIPs)-binding K268 and R270.

TULP3 functions as an adapter by connecting the IFT trafficking machinery (which it binds via its N-terminal domain, NTD: aa 1-183) to membrane proteins (which it binds via its phosphoinositide-binding C-terminal Tubby domain, CTD: aa 184-442) ^62^. To test how TULP3 binds INPP5E, we assessed co-IP of Flag-INPP5E by different EGFP-TULP3 constructs (**Fig.7b**). Flag-INPP5E specifically co-IPed with full length TULP3, an interaction that was largely dependent on TULP3’s CTD, even though weak binding to NTD was also observed (**Fig.7b**). In addition, we tested Flag-INPP5E binding to two TULP3 mutants, namely K268A+R270A and K389A. The former cannot bind phosphoinositides, and may also be hypoacetylated ^62,85^, whereas the latter removes a key lysine needed for TULP3 to interact with ARL13B and target it to cilia, according to a recent preprint ^64^. Interestingly, both mutations clearly reduced INPP5E binding, though none abolished it completely (**Fig.7b**). Therefore, our data show that INPP5E interacts with TULP3, and that this interaction is dependent on: (i) CLS2 and CLS3 in INPP5E’s catalytic domain, and (ii) TULP3’s Tubby domain and its ability to bind ARL13B and phosphoinositides (**Fig.7c**).

### INPP5E-CEP164 interaction is downregulated by CLS2-3

INPP5E also interacts with CEP164, a ciliary base protein essential for ciliogenesis. In CEP164-silenced non-ciliated cells, INPP5E fails to accumulate at the centrosome ^18^. This suggests that CEP164, by recruiting INPP5E to the ciliary base, may contribute to its ciliary targeting. Because of this, we also examined the INPP5E-CEP164 interaction. To do this, we looked at how EGFP-INPP5E and its mutants co-IP endogenous CEP164 in HEK293T cells. Interestingly, while WT, ΔCLS1, ΔCLS4 and ΔCLS1+4 all co-IPed similar amounts of CEP164, the ΔCLS2 and ΔCLS3 mutants displayed a stronger interaction, suggesting that CLS2 and CLS3 downregulate CEP164 binding (**Fig.8a**). Moreover, CEP164 strongly interacted with INPP5E’s N-terminal fragment (1-283), but not with the CLS2/3-containing C-terminal one (251-644) (**Fig.8a**). Since much more CEP164 was pulled down by EGFP-INPP5E(1-283) than EGFP-INPP5E(WT), despite both fusion proteins being expressed similarly, this further indicates that CLS2-3 antagonize CEP164 binding, which is mediated by INPP5E’s N-terminal region (**Fig.8a**).

**Figure 8.**
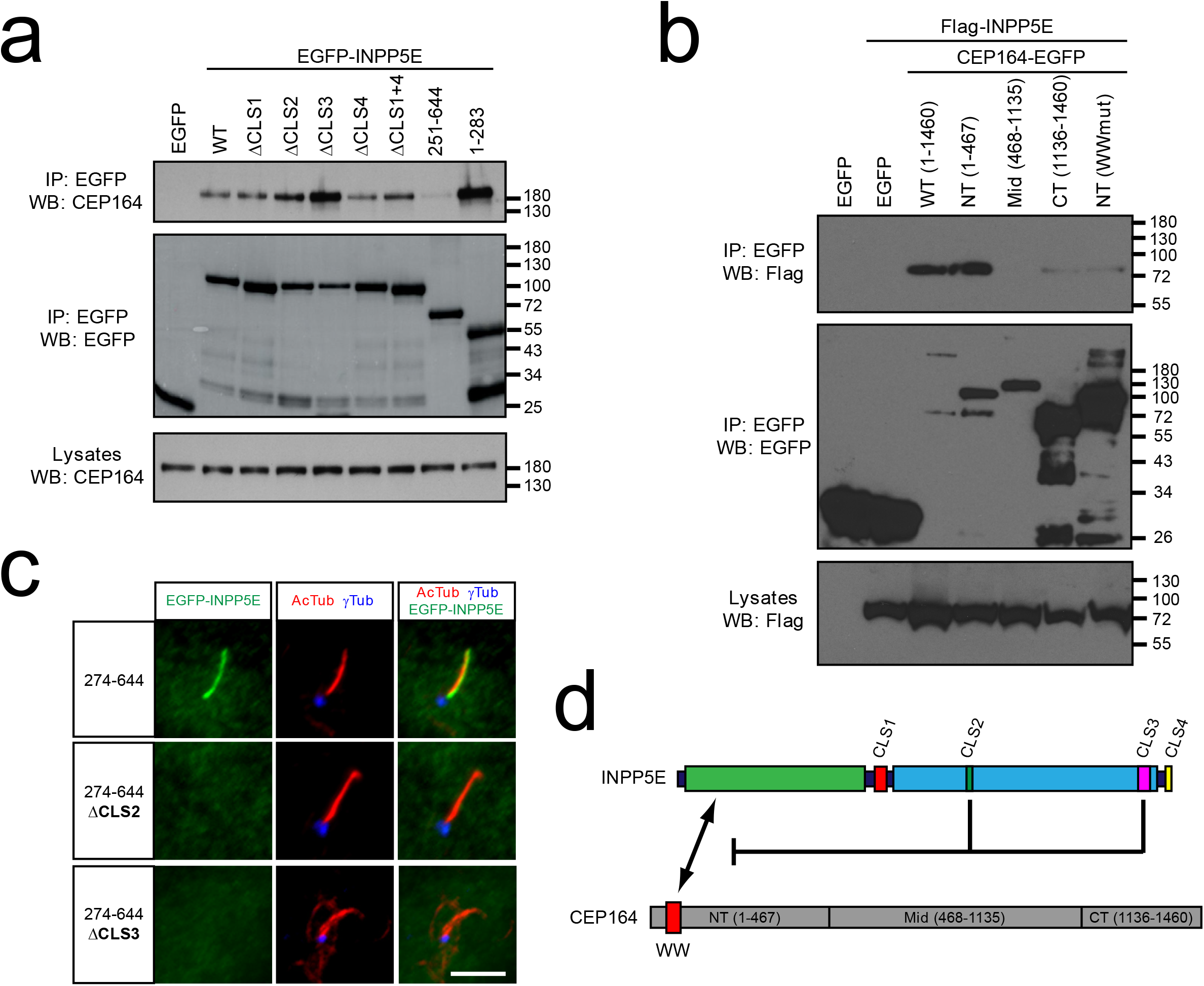
CLS2 and CLS3 antagonize INPP5E binding to CEP164 but this is not sufficient for ciliary targeting. **(a)** Lysates of HEK293T cells expressing the indicated EGFP-INPP5E variants were immunoprecipitated with GFP-Trap beads and the levels of endogenous CEP164 and exogenous EGFP were analyzed by Western blot as indicated. Molecular weight markers on the right. **(b)** Flag-INPP5E was coexpressed in HEK293T cells with the indicated CEP164-EGFP variants, including full length CEP164 (aa 1-1460), its N-terminal (NT, 1-467), middle (Mid, 468-1135) and C-terminal (CT, 1136-1460) regions, and NT carrying a mutated WW domain (WW: aa 56-89; mutation: Y74A+Y75A). Lysates were immunoprecipitated with GFP-Trap beads and analyzed by Western blot with antibodies against Flag or EGFP, as indicated. Molecular weight markers are displayed on the right. **(c)** CLS2 and CLS3 are still required for INPP5E ciliary targeting in mutants unable to bind CEP164. Cilia localization was analyzed as in previous figures for the indicated EGFP-INPP5E variants, all of which lack aa 1-273 and hence cannot bind CEP164. Scale bar, 5 μm. **(d)** Schema summarizing results from (a-b). CEP164-NT is sufficient for INPP5E binding provided the WW domain is intact. On INPP5E’s side, the proline-rich N-terminal region (aa 1-283) is sufficient to interact with CEP164. Moreover, INPP5E(1-283), INPP5E(ΔCLS2) and INPP5E(ΔCLS3) mutants all bind CEP164 more intensely than INPP5E(WT), indicating that INPP5E’s C-terminal region downregulates CEP164 binding in a CLS2/3-dependent manner. This may or may not be necessary for INPP5E ciliary targeting, but it is clearly not sufficient, as shown by the data in (c).

CEP164 contains a WW domain near its N-terminus and several coiled coils in the rest of its long sequence (1460 aa) ^65–67^. Since WW domains, like SH3 domains, interact with proline-rich ligands, we hypothesized that CEP164’s WW might interact with INPP5E’s proline-rich N-terminus ^86^. To test this, we first checked whether Flag-INPP5E was co-IPed by three CEP164 fragments spanning its N-terminal (NT, aa 1-467), middle (Mid, aa 468-1135) and C-terminal (CT, aa 1136-1460) regions ^65^. Consistent with our hypothesis, CEP164(NT)-EGFP strongly interacted with Flag-INPP5E, as did full length CEP164-EGFP (**Fig.8b**). Instead, no interaction was seen with CEP164(Mid)-EGFP and only a very weak one with CEP164(CT)-EGFP. Moreover, a mutation disrupting CEP164’s WW domain (WWmut: Y74A+Y75A) abolished INPP5E binding to CEP164(NT)-EGFP (**Fig.8b**) ^65^. Hence, CEP164’s NT is sufficient for INPP5E binding, and CEP164’s WW domain is required for it.

Altogether, these data suggest a model of CLS2-3 action: presumably, excessively strong binding to CEP164 would retain INPP5E at the ciliary base and prevent its translocation into the ciliary compartment. CLS2-3 might overcome this by loosening the CEP164-INPP5E interaction. If this is the main reason why CLS2-3 are required for INPP5E ciliary targeting, then deletion of the CEP164-interacting N-terminal region should rescue ciliary targeting of INPP5E-ΔCLS2 and INPP5E-ΔCLS3, as CLS2-3 would no longer be needed for CEP164 dissociation. To test this, we combined the ΔCLS2 and ΔCLS3 mutations with the Δ1-273 deletion, thereby generating the 274-644(ΔCLS2) and 274-644(ΔCLS3) mutants. Unlike the 274-644 control, which readily accumulated in cilia, both 274-644(ΔCLS2) and 274-644(ΔCLS3) completely failed to accumulate in cilia, just as the single ΔCLS2 and ΔCLS3 mutants (**Fig.8c**). Hence, even though CLS2-3 promote CEP164 dissociation, this is not sufficient for INPP5E ciliary targeting (**Fig.8d**). This might be due to CLS2-3 being required for binding to other ciliary trafficking proteins, such as TULP3 (**Fig.7**).

### CSNK2A1 regulates INPP5E ciliary targeting without detectable physical interaction

INPP5E prevents ectopic ciliogenesis by regulating how CEP164 interacts with TTBK2, a key ciliogenic kinase ^39^. TTBK2 was recently found to be regulated by casein kinase 2 (CSNK2A1), so we wondered whether CSNK2A1 also regulates INPP5E and, more specifically, its ciliary targeting ^87^. To test this, we examined INPP5E localization in ciliated control and *Csnk2a1*-null mouse embryonic fibroblasts. Interestingly, INPP5E ciliary levels were significantly reduced in absence of CSNK2A1 (**Fig.S6a-b**). We then performed co-IP experiments to check if CSNK2A1-myc and EGFP-INPP5E physically interact. However, we detected no such interaction, suggesting that CSNK2A1 regulates INPP5E indirectly, or through interactions too labile to detect in this manner (**Fig.S6c**).

### INPP5E-ATG16L1 interaction is modulated by CLS1 and CLS4

Recent work shows that INPP5E ciliary targeting requires ATG16L1, an autophagy protein that forms a complex with IFT-B complex component IFT20 ^68,88^. Moreover, ATG16L1 was shown to interact with INPP5E, and with its product PI4P ^68^. We therefore assessed the CLS-dependence of the ATG16L1-INPP5E interaction. Interestingly, although the single ΔCLS1, ΔCLS2, ΔCLS3 and ΔCLS4 mutants did not noticeably alter binding between EGFP-INPP5E and Flag-ATG16L1, a reduction was observed with the ΔCLS1+4 mutant, suggesting that CLS1 and CLS4 may cooperate in ATG16L1 binding, thus mirroring their cooperation in INPP5E ciliary targeting (**Fig.9a**). Consistent with CLS1 and CLS4 partaking in the interaction, the C-terminal INPP5E fragment (251-644, containing both CLS1 and CLS4) was sufficient for binding, whereas the N-terminal fragment (1-283, containing only CLS1) interacted only weakly (**Fig.9a**). These data suggest that CLS1 and CLS4 may jointly be implicated in how ATG16L1 targets INPP5E to cilia (**Fig.9b**). On the other hand, CLS2 and CLS3 would act via CEP164, TULP3 and ARL13B, with the latter also acting via CLS4, like PDE6D and RPGR (**Fig.9b**).

**Figure 9.**
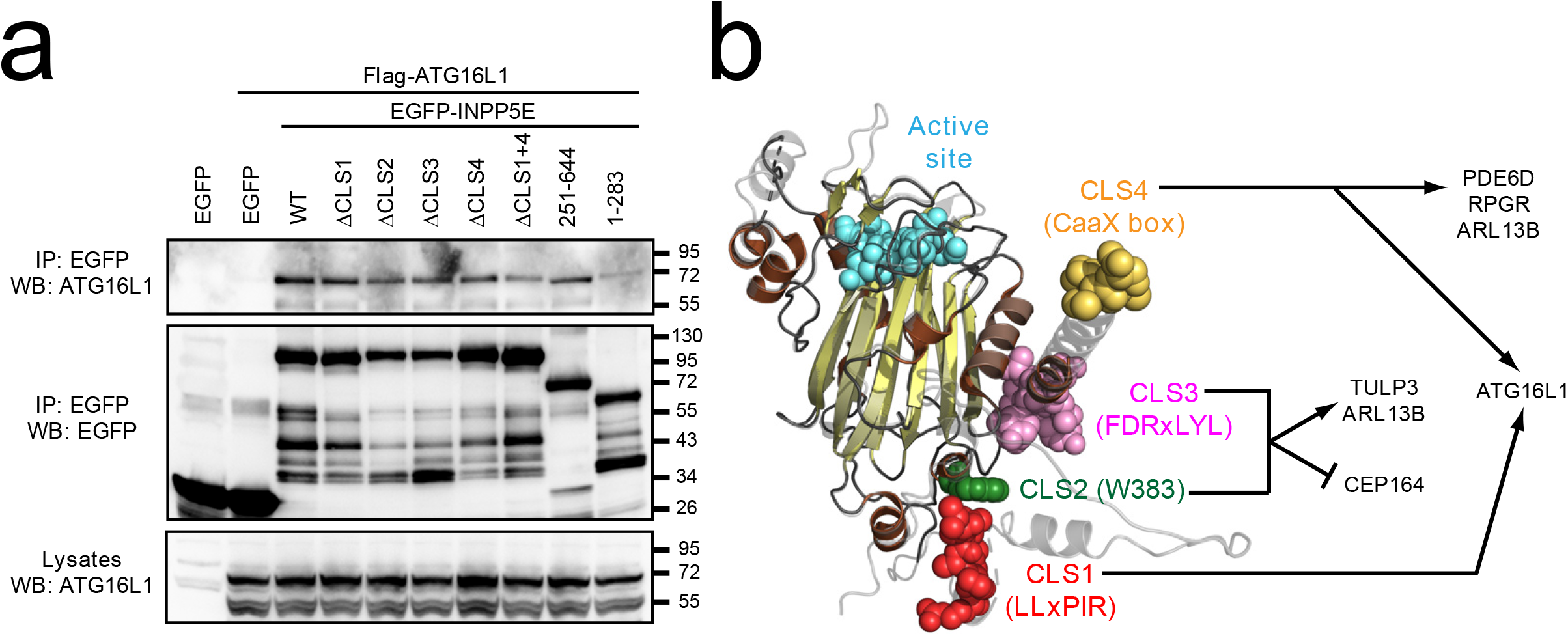
CLS1 and CLS4 modulate the ATG16L1-INPP5E interaction. **(a)** The indicated EGFP-INPP5E variants were coexpressed in HEK293T cells with Flag-ATG16L1 as indicated. Lysates were immunoprecipitated with GFP-Trap beads and analyzed by Western blot with the indicated antibodies. **(b)** Schema of INPP5E structure depicting CLS1-4 and the proteins through which they regulate INPP5E ciliary targeting, as shown herein.

### INPP5E immune synapse targeting is CLS-independent

Most cell types form primary cilia when their centrosomes are not engaged in cell division. The main exception to this is the hematopoietic lineage, where the centrosome is often engaged in other specialized structures, such as the immune synapse (IS) in lymphocytes ^89^. Interestingly, numerous parallels exist between primary cilia and the IS, such as the use of IFT trafficking machinery ^89,90^. ARL13B and ARL3 were shown to localize to the IS, and a recent preprint has shown that INPP5E does as well ^91–93^. With advice from the preprint authors, we confirmed that endogenous INPP5E accumulates at the IS between Jurkat T-cells and Raji antigen-presenting cells, a well-established IS model (**Fig.S7a-b**) ^93–96^. Given the parallels between cilia and IS, we wondered whether IS targeting of INPP5E shares the same mechanisms that INPP5E uses for ciliary targeting in other cell types. To test this, we assessed the CLS-dependence of INPP5E IS targeting. As also reported in the preprint, EGFP-INPP5E(WT) is also detected at the Jurkat-Raji IS (**Fig.S7c**) ^93^. This localization was not noticeably perturbed in the ΔCLS1, ΔCLS2, ΔCLS3, ΔCLS4 or ΔCLS1+4 mutants (**Fig.S7c**). Therefore, INPP5E IS targeting does not follow the same mechanisms as INPP5E ciliary targeting.

## DISCUSSION

In this work, we have gained many important insights into INPP5E ciliary targeting mechanisms, including among others: (i) catalytic domain integrity is essential for ciliary targeting (**Fig.1**); (ii) the FDRxLYL motif (CLS3), also essential, is part of the catalytic domain fold (**Fig.1–2**); (iii) W383 (CLS2), a solvent-exposed catalytic domain residue physically adjacent to CLS3, is specifically required for ciliary targeting (**Figs.2**, **S2-S3**); (iv) although optimal ciliary targeting requires both LLxPIR motif (CLS1) and CaaX box (CLS4), their partially redundant functions explain why none is essential for ciliary accumulation (**Fig.3**); (v) CLS1-4 are all highly conserved in vertebrates (**Fig.4**); (vi) some but not all JBTS-causative INPP5E mutations interfere with ciliary targeting (**Fig.4**); (vii) CLS4 is required for INPP5E interaction with PDE6D, ARL13B and RPGR (**Figs.5–6**, **S5**); (viii) CLS2 and CLS3 promote INPP5E binding to TULP3 via its Tubby domain (**Fig.7**); (ix) CLS2 and CLS3 downregulate binding between INPP5E and CEP164 N-termini, but this is not sufficient for ciliary targeting (**Fig.8**); (x) INPP5E ciliary targeting is regulated by CSNK2A1, with no detectable binding between them (**Fig.S6**); (xi) INPP5E binding to ATG16L1 is reduced by concomitant mutation of CLS1 and CLS4, but not by the single mutations (**Fig.9**); and (xii) INPP5E IS targeting is CLS-independent (**Fig.S7**). For a summary of how each CLS affects binding to PDE6D, RPGR, ARL13B, TULP3, CEP164 and ATG16L1, see **Table 1**.

**Table 1.** CLS-dependence of INPP5E protein-protein interactions. Cilia localization and the indicated interactions are shown for each EGFP-INPP5E construct on the left column. For both localization and interactions, meaning of arrows is as follows: two upward green arrows (strong), one upward green arrow (moderate), one downward red arrow (low), and two downward red arrows (undetectable).

Multiple CLSs Target INPP5E to Cilia
| INPP5E construct | Ciliary? | INPP5E interactors |  |  |  |  |  |
| --- | --- | --- | --- | --- | --- | --- | --- |
|  |  | PDE6D | RPGR | ARL13B | TULP3 | CEP164 | ATG16L1 |
| WT | ↑↑ | ↑ | ↑ | ↑ | ↑ | ↑ | ↑ |
| ΔCLS1 | ↑ | ↑ | ↑ | ↑ | ↑ | ↑ | ↑ |
| ΔCLS2 | ↓↓ | ↑ | ↑ | ↓ | ↓ | ↑↑ | ↑ |
| ΔCLS3 | ↓↓ | ↑ | ↑ | ↓ | ↓ | ↑↑ | ↑ |
| ΔCLS4 | ↑ | ↓↓ | ↓↓ | ↓↓ | ↑ | ↑ | ↑ |
| ΔCLS1+4 | ↓↓ | ↓↓ | ↓↓ | ↓↓ | ↑ | ↑ | ↓ |
| CT (251-644) | ↑↑ | ↑ | ↑ | ↑ | ↑ | ↓↓ | ↑ |
| NT (1-283) | ↓↓ | ↓↓ | ↓↓ | ↓↓ | ↓ | ↑↑ | ↓ |

Since catalytic domain integrity is essential for ciliary localization, it is important to distinguish whether candidate CLS residues act specifically, engaging ciliary trafficking machinery, rather than non-specifically, preserving domain structure. By using enzyme activity as a readout for catalytic domain integrity, we demonstrated that all four CLSs act specifically, since: (i) simultaneous deletion of both CLS1 and CLS4 had no effect on catalysis (**Fig.S4**); and (ii) although deleting CLS2 or CLS3 reduced activity by about 2-fold, this is insufficient to explain the virtually complete loss of ciliary targeting observed with these mutants (**Fig.2**).

Further supporting this conclusion are the following considerations: (i) an equivalent 2-fold reduction was also observed with R379A, yet this mutant was readily observed in cilia, even though R379 is directly adjacent to CLS2-3 in the folded catalytic domain (**Fig.2**); (ii) the same applies to the JBTS mutant R345S, whose activity is also 2-fold lower than WT (**Fig.4** and data not shown); (iii) as shown by the cycloheximide chase assays, over 50% of the ΔCLS2 and ΔCLS3 proteins have a long half-life, similar to WT, suggesting that these mutations only interfere with catalytic domain integrity, and hence stability, in less than half of the protein population (**Fig.S2**); (iv) W383 mutation to alanine, valine, leucine, isoleucine and methionine all led to a modest reduction in activity, yet a complete loss of ciliary targeting (**Fig.S3**); and (v) CLS2 and CLS3 specifically affect binding to some but not all INPP5E interactors (**Fig.5–9**). Therefore, we conclude that CLS1-4 are all *bona fide* CLSs.

Although mutations in other JBTS genes are known to disrupt INPP5E ciliary localization, mutations in *INPP5E* itself had not been reported to do so ^3,5,13–20^. Instead, INPP5E mutations were shown to impair INPP5E enzyme activity, without affecting ciliary targeting ^8,29^. Here, we show for the first time that *INPP5E* gene mutations do sometimes prevent ciliary accumulation. Curiously, the mutations causing such mistargeting were not the ones located near CLS2-3, but rather mutations that likely compromise catalytic domain integrity (**Fig.4**). Still, how G286R, V303M and W474R disrupt ciliary targeting has not been addressed empirically. Interestingly, both G286R and W474R are homozygous mutations, suggesting a complete lack of INPP5E in the cilia of these patients. Whether and how this contributed to JBTS manifestations in these patients, as compared to patients with activity-impaired INPP5E in their cilia, remains an open question.

Besides discovering two novel CLSs in INPP5E (CLS1-2), we also shed light into how these novel CLSs relate to the previously identified ones (CLS3-4). For instance, we unexpectedly found that CLS1 and CLS4, despite being far apart from each other, cooperatively promote ciliary targeting. Although single CLS1 and CLS4 mutants show moderately impaired cilia localization, it is only in the double mutant that ciliary targeting is completely abolished (**Fig.3a-c**). This indicates a partial functional redundancy that begs for an explanation. Although the mechanism of CLS4 action is fairly well understood, how CLS1 can partially substitute for it remains unclear ^17,50,51^. In this regard, we tested several hypotheses. First, we hypothesized that CLS1, like CLS4, might also promote interaction with PDE6D, but the answer was clearly negative (**Fig.5**, **Fig.9b**). We then wondered whether CLS1 might act via RPGR, a prenylated INPP5E and PDE6D-interacting protein also needed for INPP5E ciliary targeting ^54–59,97^. Again, however, we found that CLS4 but not CLS1 mediates INPP5E-RPGR binding (**Fig.S5**, **Fig.9b**). ARL13B and ARL3 promote dissociation of the PDE6D-INPP5E complex inside cilia. Nevertheless, ARL3 does not interact with INPP5E ^18^ (data not shown), whereas ARL13B binding also depends on CLS4 but not CLS1 (**Fig.6**, **Fig.9b**). For TULP3 and CEP164, neither CLS1 nor CLS4 had any effect on the interactions (**Figs.7–8**, **Fig.9b**).

We also looked at ATG16L1, an autophagy protein without which INPP5E cannot accumulate in cilia ^68^. As previously reported, we detected INPP5E-ATG16L1 binding, which was reduced with ΔCLS1+4 but not ΔCLS1 or ΔCLS4 (**Fig.9a**). This mirrors the effects of CLS1 and CLS4 on ciliary targeting, suggesting that ATG16L1 may exert its effects via these CLSs (**Fig.9b**). Since ATG16L1 is involved in the Golgi exit of IFT20, an IFT-B component that traffics from Golgi to cilia base, one could speculate that ATG16L1 also promotes INPP5E Golgi exit ^68^. This would be in accordance with previous reports of INPP5E Golgi localization, and with post-prenylation processing of CaaX box proteins occurring on the ER-Golgi surface ^11,79^. However, how ATG16L1 promotes INPP5E ciliary accumulation, and to what extent ATG16L1 binding explains the combined CLS1+CLS4 effects remains unknown.

Besides the functional CLS1-CLS4 connection, we also uncovered a CLS2-CLS3 link. Given their steric proximity (**Fig.2b**), we hypothesize CLS2 and CLS3 work together as a functional unit to recruit ciliary trafficking proteins like ARL13B and TULP3 (**Figs.6–7**, **Fig.9b**). Consistent with this, CLS2 and CLS3 behaved largely equivalently in all our interaction experiments (**Figs.5–9**). Moreover, the need to keep CLS2-3 together also explains why a folded catalytic domain is critical for ciliary targeting (**Fig.1**). Future structural studies should further clarify these issues.

ARL13B regulates INPP5E ciliary targeting in at least two ways: (i) by directly interacting with INPP5E in a CLS3-dependent manner; and (ii) by functioning as an ARL3-GEF, thereby promoting INPP5E release from PDE6D inside cilia ^17,18,48–50^. However, of these two regulatory modes, only the direct interaction is essential for ARL13B to mediate INPP5E ciliary retention ^48,49^. Our ARL13B-INPP5E interaction experiments showed that CLS2, CLS3 and CLS4 all affect their association (**Fig.6**). Given that CLS2-3 likely form a functional unit as mentioned above, CLS2 probably affects the direct ARL13B-INPP5E interaction, as previously reported for CLS3 ^18^. CLS4 mutation has also been shown recently to moderately reduce ARL13B binding ^48^. A possible explanation is that, by mediating INPP5E membrane insertion, farnesylation facilitates access to ARL13B, a fatty acylated protein ^98,99^.

TULP3 is essential for ciliary targeting of both ARL13B and INPP5E, among other ciliary membrane proteins ^61,64^. Through its C-terminal Tubby domain, TULP3 directly interacts with ARL13B, targeting it to cilia, which in turn allows INPP5E ciliary targeting ^64^. However, whether TULP3 and INPP5E interact with each other had not been reported. Using co-IPs in HEK293T cells, we show that this is indeed the case, with the interaction depending on CLS2 and CLS3 (**Fig.7a**). On TULP3’s side, the interaction mostly depends on the Tubby domain, with a minor contribution from the NTD (**Fig.7b**). Moreover, Tubby domain mutations K389A and K268A+R270A also reduced binding, more so with K389A (**Fig.7b**). The K268A+R270A mutation interferes with TULP3’s ability to bind phosphoinositides and target GPCRs and other transmembrane proteins to cilia, but has no effect on TULP3’s ability to target ARL13B and INPP5E to cilia ^64^. Thus, K268 and R270, despite moderately affecting TULP3-INPP5E binding, do not seem to play an important role in INPP5E ciliary accumulation. On the other hand, K389 is important for ARL13B binding and critical for its ciliary targeting and that of INPP5E ^64^. This suggests that the TULP3-INPP5E interaction may be mediated by ARL13B. Consistently, the K389-dependent TULP3-ARL13B interaction is known to be direct, whereas the TULP3-INPP5E interaction in our co-IPs could well be indirect ^64^. As a counterargument, if INPP5E can only bind TULP3 indirectly via ARL13B, then it is hard to explain why mutation of CLS4 reduces INPP5E-ARL13B but not INPP5E-TULP3 binding (**Figs.6–7**). Clearly more experiments are needed to clarify these points.

CEP164 binds INPP5E and recruits it to the ciliary base ^18^. Herein, we show that this interaction involves the N-termini of both proteins, with the WW domain in CEP164 being critical (**Fig.8a-b**). Since WW domains typically interact with proline-rich motifs, which abound in INPP5E’s N-terminal region, it seems very likely that the CEP164-INPP5E interaction also involves these motifs ^86^. In addition to defining this core interaction, we also provide evidence that this interaction is negatively regulated by CLS2-3 in INPP5E’s catalytic domain (**Fig.8a**). This is an interesting observation, as the CEP164-INPP5E interaction may need to be loosened for INPP5E to efficiently enter cilia. Such loosening, however, cannot be the main function of CLS2-3, as they were still required for ciliary targeting of INPP5E mutants unable to bind CEP164 (**Fig.8c**).

Tau tubulin kinase 2 (TTBK2) is an essential ciliogenic kinase that, like INPP5E, binds CEP164’s WW domain via a proline-rich region ^65,100^. Thus, INPP5E and TTBK2 may compete for CEP164 binding. Moreover, INPP5E’s product PI4P binds both CEP164 and TTBK2, blocking their interaction ^39^. These mechanisms help explain why INPP5E antagonizes ectopic ciliogenesis in non-ciliated cells ^39^. On the other hand, these mechanisms suggest that INPP5E targeting and function may be regulated by TTBK2. Given that TTBK2-null cells fail to form cilia, we assessed this possibility in another model: in cells lacking the casein kinase CSNK2A1, a recently reported upstream regulator of TTBK2 ^87^. Indeed, CSNK2A1-KO cells displayed reduced ciliary INPP5E levels (**Fig.S6a**). However, since CSNK2A1 has multiple substrates, whether this effect is TTBK2-dependent remains uncertain. We also looked for a CSNK2A1-INPP5E interaction but found none, suggesting CSNK2A1’s effects on INPP5E are indirect or mediated by interactions that are too weak or transient for us to detect (**Fig.S6c**).

Finally, given the known parallels between primary cilia and immune synapses, with the latter containing, among other ciliary proteins, ARL13B and INPP5E, we asked whether CLS1-4 also drive INPP5E targeting to the IS ^92,93^. The answer was no, pointing to clearly distinct mechanisms for targeting to both structures (**Fig.S7**). This may mean that INPP5E targeting to cilia and IS evolved independently of each other or, alternatively, that a common evolutionary origin has been blurred by a long history of evolutionary divergence. The fact that INPP5E IS targeting is quickly induced upon assembly of the highly plastic and dynamic IS also suggests a more transient role for INPP5E at the IS, as opposed to its more constitutive ciliary localization. A deeper knowledge of how INPP5E is targeted and functions at these signaling platforms, cilia and IS, will be needed to answer these questions.

Altogether, our data show that INPP5E ciliary targeting is a surprisingly complex process involving four different cis-acting sequences (CLS1-4), and multiple trans-acting factors (like PDE6D, RPGR, ARL13B, TULP3, CEP164 and ATG16L1). This level of complexity is unusual, especially when compared to other ciliary cargoes, whose targeting is more straightforward, typically involving a single CLS ^1,70–73^. The complexity and redundancy in INPP5E ciliary targeting suggest this is a very important process, subject to fine regulation. This is consistent with the surprisingly wide range of functions INPP5E plays at the cilium, controlling among others their lipid and protein composition, assembly and disassembly, exovesicle release, and signaling ^7,8,12,32–45^. By providing a much broader view of the mechanisms involved in INPP5E ciliary targeting, the stage is now set for a deeper molecular understanding of these processes and their regulation.

## METHODS

### Antibodies and reagents

Mouse monoclonal antibodies: anti-acetylated α-tubulin (AcTub) (Sigma, T7451, clone 6-11B-1, IF: 1:10,000), anti-α-tubulin (Proteintech, 66031-1-Ig, WB: 1:1000), anti-EGFP (Proteintech, 50430-2-AP, WB: 1:1000), anti-ARL13B (Proteintech, 66739-1-Ig, WB: 1:1000), anti-Flag (Sigma, F3165, clone M2, WB: 1:2000). Rabbit polyclonal antibodies: anti-EGFP (Proteintech, 50403-2-AP, IF: 1:200, WB: 1:1000), anti-RPGR (Proteintech, 16891-1-AP, WB: 1:1000), anti-TULP3 (Proteintech, 13637-1-AP, WB: 1:2000), anti-CEP164 (Proteintech, 22227-1-AP, WB: 1:1000), anti-ATG16L (MBL, PM040, WB: 1:1000), anti-Myc (Proteintech, 16286-1-AP, WB: 1:1000), and anti-INPP5E (Proteintech, 17797-1-AP, IF: 1:100). Goat polyclonal antibody: anti-γ-tubulin (γ-Tub) (Santa Cruz, sc-7396, IF: 1:200). AlexaFluor (AF)-conjugated donkey secondary antibodies for IF (from Thermofisher, used at 1:10,000): AF488 anti-rabbit IgG (A21206), AF555 anti-mouse IgG (A31570) and AF647 anti-goat IgG (A21447). Also from Thermofisher were HRP-conjugated goat anti-mouse/rabbit IgG secondary antibodies for WB (all at 62ng/ml) and AF546 Phalloidin (A22283), used at 1:100. For IP assays, GFP-Trap_MA beads (Chromotek, gtma-20) were used. For phosphatase assays, PtdIns(4,5)P2-diC8 (P-4508) and Malachite Green Assay Kit (K-1500) were from Echelon Biosciences (Tebu-Bio), while n-octyl-β-D-glucopyranoside was from Alfa Aesar (J67390.03).

### Plasmids and mutagenesis

Plasmids encoding human INPP5E-WT and D477N in the pEGFP-C1 backbone (pEGFP-INPP5E-WT and D477N) were reported previously ^7,32^. All other EGFP-INPP5E mutants were created by site-directed mutagenesis using overlap extension PCR (or single-step PCR where possible) to amplify the mutagenized ORFs and clone them into pEGFP-C1 using XhoI-KpnI sites. Amplifications were performed with Platinum SuperFi DNA polymerase (Thermofisher), and all finished constructs were validated by DNA sequencing (Eurofins Genomics). To obtain pFlag-INPP5E, the INPP5E ORF was cloned into the EcoRI-KpnI sites of pFlag-CMV4 vector. To generate pFlag-PDE6D, human PDE6D ORF was amplified with primers encoding N-terminal Flag, and Flag-PDE6D was then used to replace the AgeI-EcoRI-flanked mVenus-PDE6D ORF in the mVenus-PDE6D plasmid described below.

Plasmid pcDNA3.1(+)-N-DYK-RPGR, expressing Flag-tagged human RPGR (NM_000328.3), was purchased from GenScript. To generate pEGFP-TULP3(K389A), overlap extension PCR was used to obtain the mutant ORF, which was then cloned KpnI-BamHI into pEGFP-C1. All other TULP3-expressing plasmids were described elsewhere ^101^. CEP164-EGFP was from Addgene (#41149), and was mutagenized to obtain the NT, NT-WWmut, Mid and CT versions ^65^. To generate pFlag-ATG16L1, EcoRI insert from pMRX-IP-SECFP-hATG16A1 (Addgene #58994) was transferred to pFlagCMV4. FKBP ORF was cloned into p-mCherry-C1 to obtain mCherry-FKBP, into which INPP5E and its mutants were subcloned. mVenus-PDE6D and ARL13B-EYFP were obtained by inserting the human ORFs into p-mVenus-C1 and pEYFP-N1, respectively. FRB-CFP-CaaX box is described elsewhere ^102^.

### Cell culture and transfections

All cell lines were grown at 37 °C and 5% CO2 in a humidified atmosphere and were regularly tested to ensure they were mycoplasma-free. hTERT-RPE1 cells were cultured in DMEM/F12 basal medium supplemented with 10% fetal bovine serum (FBS) and were reverse transfected using JetPrime (Polyplus-transfection), and their cilia analyzed 48 hours later, after 24 hours of serum starvation. HEK293T cells were maintained in DMEM+10% FBS, transfected using the calcium phosphate method, and lysed 40-48 hours later. HeLa cells were cultured in DMEM+10% FBS+Penicillin/Streptomycin and transfected using FuGENE 6 (Promega). Co-recruitment assays were performed 24 hours after transfection. CRISPR-engineered control and *Csnk2a1*-null mouse embryonic fibroblasts have been described elsewhere ^87^. Raji B and Jurkat T (clone JE6.1) cell lines from ATCC were cultured in RPMI-1640 medium containing L-glutamine, penicillin/streptomycin, and 10% heat-inactivated FBS.

### Immunofluorescence microscopy

hTERT-RPE1 cells grown to confluence on coverslips were fixed 5 min in PBS+4% paraformaldehyde (PFA) at room temperature (RT), followed by freezer-cold methanol for 3 min at −20°C. Cells were then blocked and permeabilized for 30-60 min at RT in PBS+0.1% Triton X100+2% donkey serum+0.02% sodium azide (blocking solution). Coverslips were then incubated in a humidified chamber for 2 hours at RT (or overnight, 4°C) with blocking solution-diluted primary antibodies. After three PBS washes, PBS-diluted secondary antibodies and DAPI (1 μg/ml, Thermofisher) were added for 1 hour at RT in the dark. After three more PBS washes, coverslips were mounted on slides using Prolong Diamond (Thermofisher), incubated overnight at 4°C, and imaged with a Nikon Ti fluorescence microscope. Brightness and contrast of microscopic images were adjusted for optimal visualization using Adobe Photoshop or Fiji (Image J).

### Immunoprecipitation and Western blot

HEK293T cells were lysed 40-48 hours post-transfection in buffer containing 50mM Tris-HCl pH=7.5, 150mM NaCl, 1% Igepal CA-630 (Sigma-Aldrich) and 1X Halt protease inhibitor cocktail (Thermofisher, #78429). Lysates were then rotated (15 min, 4°C) and centrifuged (10 min, 20,000g, 4°C), and protein levels in the postnuclear supernatants measured with Pierce BCA Protein Assay Kit (Thermofisher). After equalizing protein amount and concentration in all samples, EGFP fusion proteins were immunoprecipitated with GFP-Trap magnetic agarose (GFP-Trap_MA, Chromotek) beads for 2 hours or overnight at 4°C with rotation. Beads were then washed thrice in lysis buffer without protease inhibitors, eluted with 2X Laemmli buffer containing 200mM DTT, and boiled 5 min at 95°C. SDS-PAGE and Western blots were performed as previously described ^101,103^. For activity assays using immunoprecipitates of EGFP-INPP5E fusion proteins, see next section.

### Phosphoinositide phosphatase assays

Activity assays were performed essentially as described ^8^. Briefly, HEK293T cell lysates were obtained as above, and their protein levels measured with Pierce BCA Protein Assay kit (Thermofisher). EGFP-INPP5E or its mutants were then immunoprecipitated from 1 mg of cell lysate by overnight rotation at 4°C with GFP-Trap_MA beads (Chromotek). Beads were then washed thrice in buffer containing 50mM Tris-HCl pH=7.5 and 150mM NaCl buffer (no protease inhibitors or detergent), and twice more in activity buffer (50mM Tris-HCl pH=7.5, 150mM NaCl, 3mM MgCl2 and 0.1% octyl-β-D-glucopyranoside (Alfa Aesar)). For the activity assays, beads were incubated in activity buffer supplemented with 120 μM diC8-PtdIns(4,5)P2, from Echelon Biosciences. After incubating the enzyme reactions for 20 min at 37°C, the supernatant was retrieved from the beads and its phosphate concentration measured at 620 nm using the Malachite Green Assay Kit (Echelon Biosciences). Beads were then processed for Western blot as above.

### Co-recruitment assays

Co-recruitment assays were performed using Eclipse Ti microscope (Nikon, Japan) with a 100X TIRF objective (1.0X zoom and 4X4 binning) in TIRF mode and PCO-Edge 4.2 BI sCMOS camera (PCO, Germany), driven by NIS Elements software (Nikon) and equipped with 440 nm, 514 nm and 561 nm laser lines. Time lapse imaging was performed at 2 min intervals for 20 minutes, with 100 nM rapamycin addition after the fifth time point. All live cell imaging was conducted at 37°C, 5% CO2 and 90% humidity with a stage top incubation system (Tokai Hit). Vitamin and phenol red-free media (US Biological) supplemented with 2% FBS were used in imaging to reduce background and photobleaching. Adequate co-expression of all relevant plasmids was confirmed by fluorescent imaging at appropriate wavelengths. To minimize variability due to relative expression levels, only cells showing at least 30% increase in mCherry intensity after addition of rapamycin were considered for quantification. All image processing and analyses for co-recruitment assays were performed using Metamorph (Molecular Devices, Sunnyvale, CA, USA) and FIJI software (NIH, Bethesda, MD, USA).

### Immune synapse analyses

Raji cells were attached to glass-bottom microwell culture dishes (IBIDI) using poly-L-lysine (20 μg/mL). Raji cells were then labeled with 10 μM CMAC (7-amino-4-chloromethylcoumarin, Molecular Probes), pulsed with 1 μg/ml SEE (Staphylococcus enterotoxin E, Toxin Technologies), and mixed with Jurkat T cells (clone JE6.1) (ATCC). To promote synaptic conjugate formation, cell-containing dishes were centrifuged at low speed (200xg, 30 seg) and incubated 5 min at 37°C. For recombinant protein expression, exponentially growing Jurkat T cells were electroporated with 20–30 μg of plasmid as previously reported ^94^, and Raji cells were added as above 40-48 hours post-electroporation. Cell conjugates were then fixed, first with PBS+2% PFA (10 min, RT), then with cold acetone (10 min at −20°C). Immunofluorescence staining was done as previously described ^94^. Imaging was performed using a Nikon Eclipse TiE microscope equipped with a DS-Qi1MC digital camera, a PlanApo VC 60x NA 1.4 objective, and NIS-AR software (all from Nikon). Epifluorescence images were then deconvolved with Huygens Deconvolution Software from Scientific Volume Image (SVI), using the “widefield” optical option.

### Structural analyses

Swiss PDB Viewer was used to visualize the crystallographic structure of INPP5E aa 282-623 (Protein Data Bank (PDB) ID: 2xsw) in order to identify candidate CLS or ciliopathy residues near the FDRxLYL motif ^75^. Final figures were rendered using the PyMOL Molecular Graphics System (Version 2.0 Schrödinger), from either the PDB structure or the full length AlphaFold model (AF-Q9NRR6-F1) ^76^.

## Statistical analyses

Graphs and statistical analyses were created using GraphPad Prism 8 software. Activity assays were analyzed by one-way ANOVA, followed by Tukey’s multiple comparisons tests. The corresponding figure legends contain the specific details of each experiment. Co-recruitment assay graphs show means ± 95% confidence interval with n≥ 40 different cells pooled from at least three independent experiments.

## AUTHOR CONTRIBUTIONS

DCR, RMM and FRGG conceived and designed the study. DCR and RMM performed most experiments. PBG, ADR, AL, MBSR and GH also performed experiments. DCR, RMM, PBG, ADR, AL, OP, SG, MR, MI, TI and FRGG conceptualized experiments and analyzed data. SG, MI, TI and FRGG supervised the work. FRGG wrote the manuscript, with input from all other authors. All authors read and agreed to the published version of the manuscript.

## ACKNOWLEDGEMENTS

We thank Dr. Jung-Chi Liao for advice with INPP5E immune synapse stainings. This publication is part of grants PID2019-104941RB-I00 (to FRGG) and PID2020-114148RB-I00 (to MI), both funded by MCIN/AEI/ 10.13039/501100011033, and the latter also funded by ERDF, a way of making Europe. FRGG is also recipient of an ACCI-2020 grant from CIBERER. This work was also funded by American Heart Association fellowship 20POST35220046 (ADR), discretionary funds (TI), Community of Madrid contract PEJD-2016/BMD-2341 (MBSR), and MICINN predoctoral grant BES2016-077828 (RMM).

## SUPPLEMENTARY MATERIAL

The Supplementary Material for this article includes Supplementary Figures S1-S7 and their legends.

