## Supplemental Figures S1-S7 for "Multiple Ciliary Localization Signals Control INPP5E Ciliary Targeting"

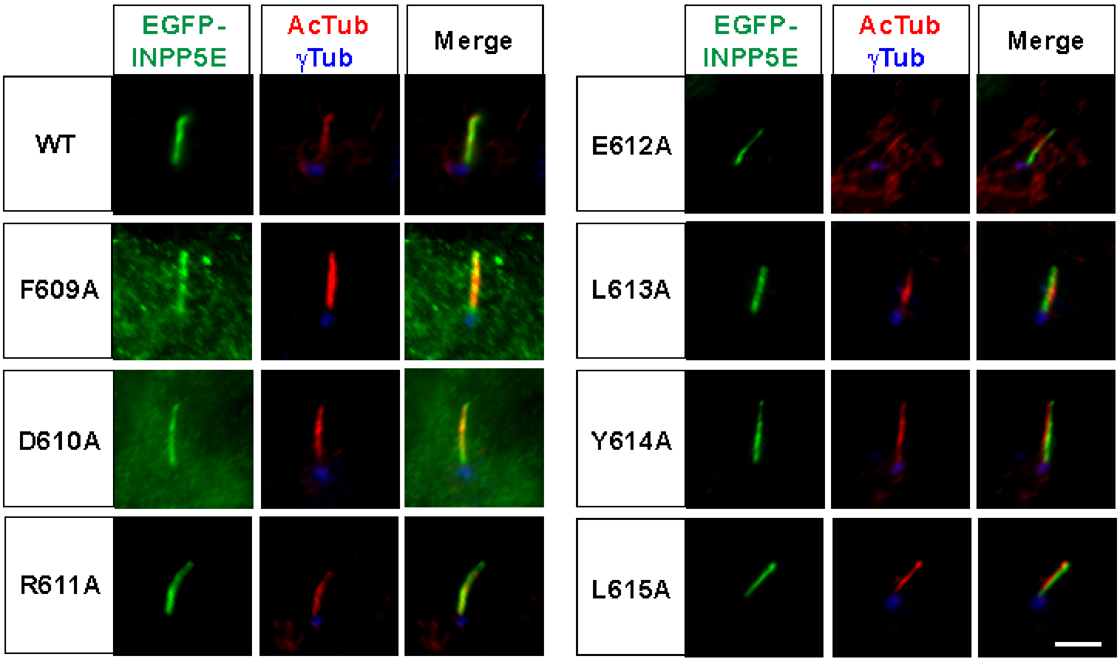


**Figure S1. FDRELYL motif residues are not individually required for INPP5E cilia localization.** Ciliary targeting of EGFP-INPP5E wild type (WT) or the indicated FDRELYL motif mutants was assessed by immunofluorescence microscopy in transfected hTERT-RPE1 cells, which were stained with antibodies against acetylated α-tubulin (AcTub), γ-tubulin (γTub) and EGFP to detect the fusion proteins. Scale bar, 5 µm.

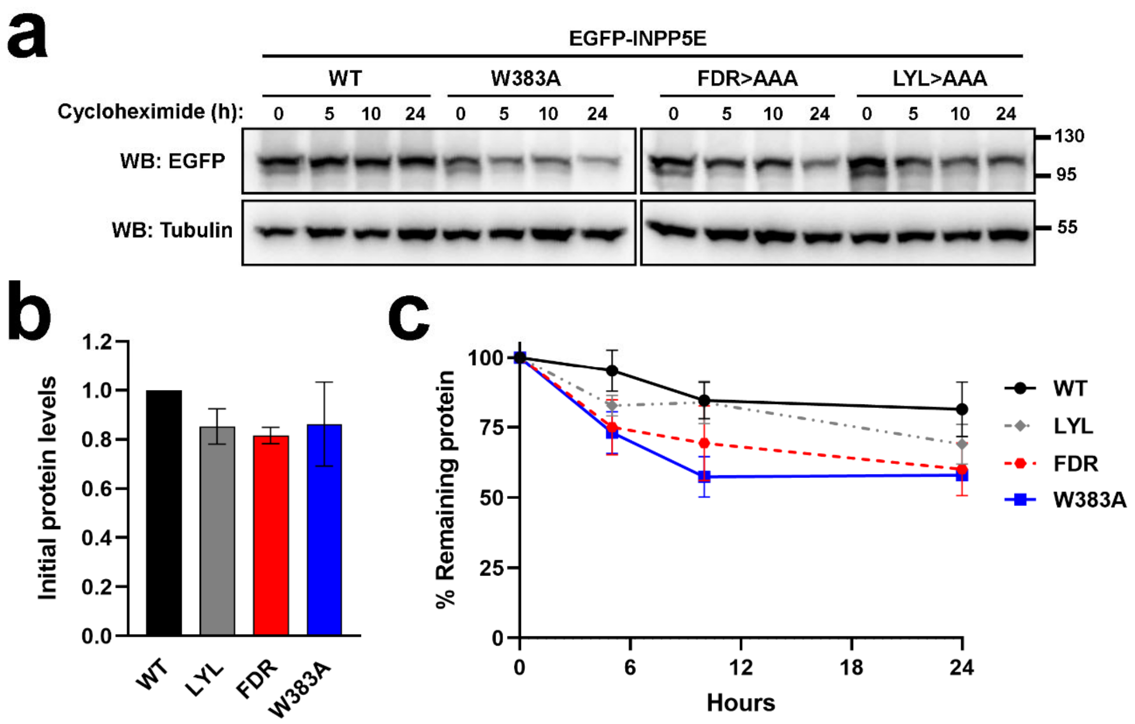


**Figure S2. Stability of W383A and FDRxLYL mutants. (a)** The indicated EGFP-INPP5E forms were expressed in HEK293T cells for 24 hours, at which time 200µg/ml cycloheximide was added for the indicated number of hours. Cell lysates were then analyzed by Western blot with antibodies against EGFP and α-tubulin. Molecular weight markers in kDa shown on the right. FDR>AAA: F609A+D610A+R611A; LYL>AAA: L613A+Y614A+L615A. **(b)** Protein levels at the time of cycloheximide addition for the same EGFP-INPP5E forms as in (a). EGFP/Tubulin band intensity ratios were normalized to WT and plotted as mean±SEM from n=3 independent experiments, including one in (a). One-way ANOVA revealed no significant differences. **(c)** Time course of protein levels after cycloheximide addition. Amounts are EGFP/Tubulin ratios for each EGFP-INPP5E form, normalized to the ratio at 0 hours. Data are mean±SEM (n=3 independent experiments). Unpaired t-tests reveal non-significance except for WT versus W383A at 10 hours (p=0.048).

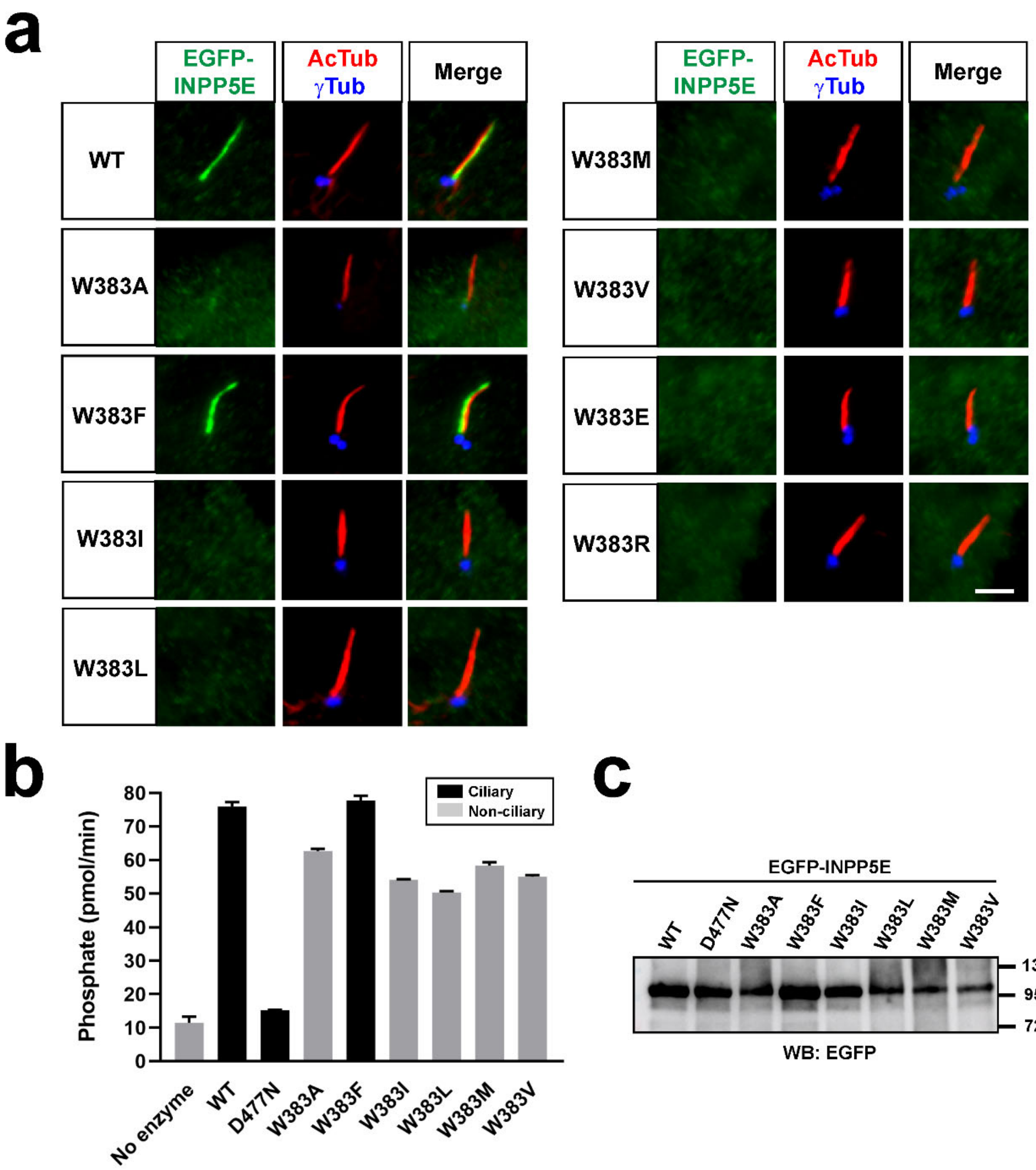


**Figure S3. Effect of W383 mutations on ciliary targeting and activity. (a)** Cilia localization of the indicated EGFP-INPP5E variants was analyzed in hTERT-RPE1 cells as in Figures 1-2. Scale bars, 5 µm. **(b)** 5-phosphatase activity, expressed as picomoles of released inorganic phosphate per minute, was measured, using PI(4,5)P2 as substrate, in immunoprecipitates of HEK293T cells transfected with the indicated EGFP-INPP5E variants. Cilia-localized constructs shown as black columns, non-ciliary as grey. Data are mean±SEM of n=3 technical replicates. **(c)** Western blot of the anti-EGFP immunoprecipitates used for the activity assays in (b). Molecular weight markers are on the right (kDa).

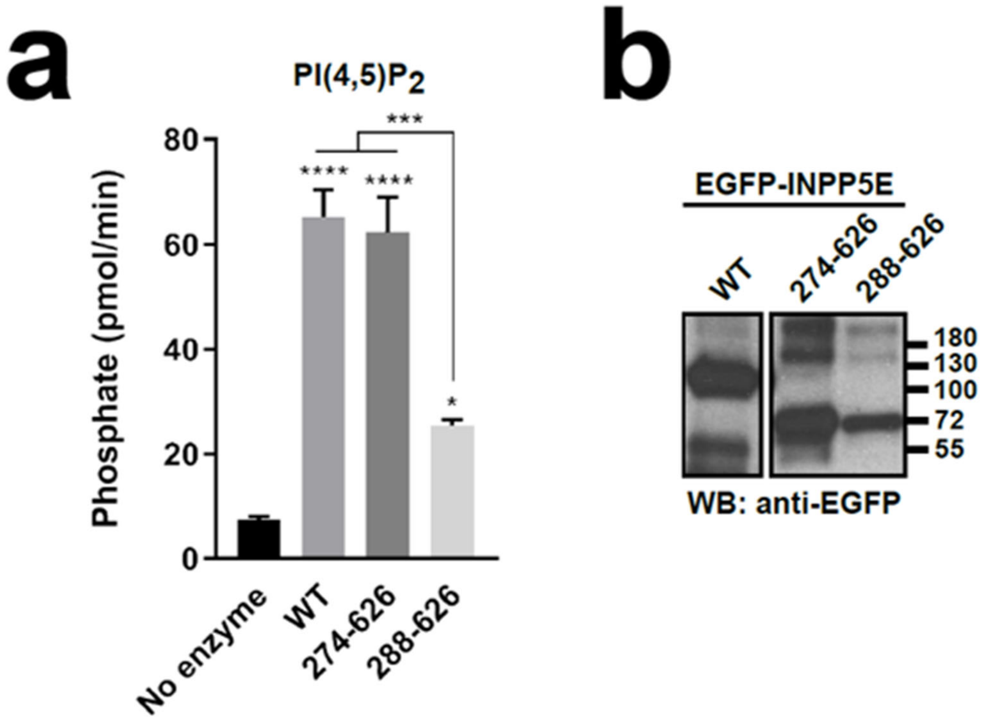


**Figure S4. CaaX box and the CLS-containing residues 251-273 do not affect enzyme activity. (a)** Phosphatase activity, expressed as picomoles of released inorganic phosphate per minute, was measured in immunoprecipitates of HEK293T cells transfected with the indicated EGFP-INPP5E variants, or in control buffer (no enzyme). Activity was measured using PI(4,5)P2 as substrate. Data are shown as mean±SEM of n=3 independent experiments. Data were analyzed by one-way ANOVA followed by Tukey's multiple comparisons tests. Significance (relative to no enzyme unless otherwise indicated) shown as $p<0.05$(*), $p<0.001$(***) or $p<0.0001$(****). **(b)** Representative anti-EGFP immunoblot showing protein levels of EGFP-INPP5E variants in the immunoprecipitates used in (a). All three samples were run in the same SDS-PAGE gel and immunoblotted and detected in parallel. Molecular weight markers, in kDa, are shown on the right.

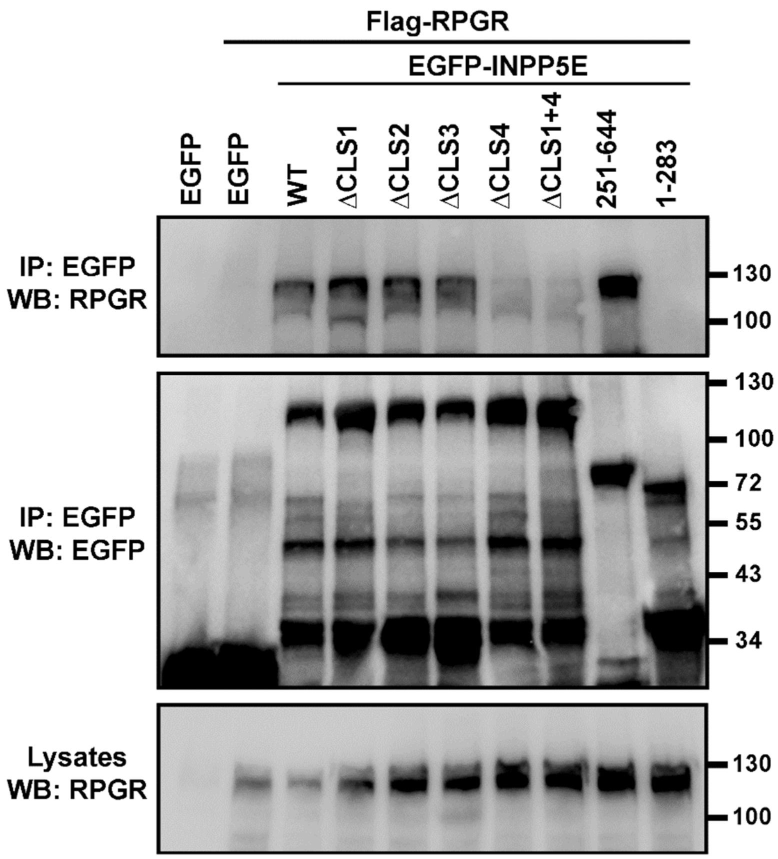


**Figure S5. CLS4 promotes INPP5E binding to RPGR.** The indicated EGFP-INPP5E variants were coexpressed in HEK293T cells with Flag-RPGR, as indicated. Lysates were immunoprecipitated with GFP-Trap beads and analyzed by Western blot with the indicated antibodies.

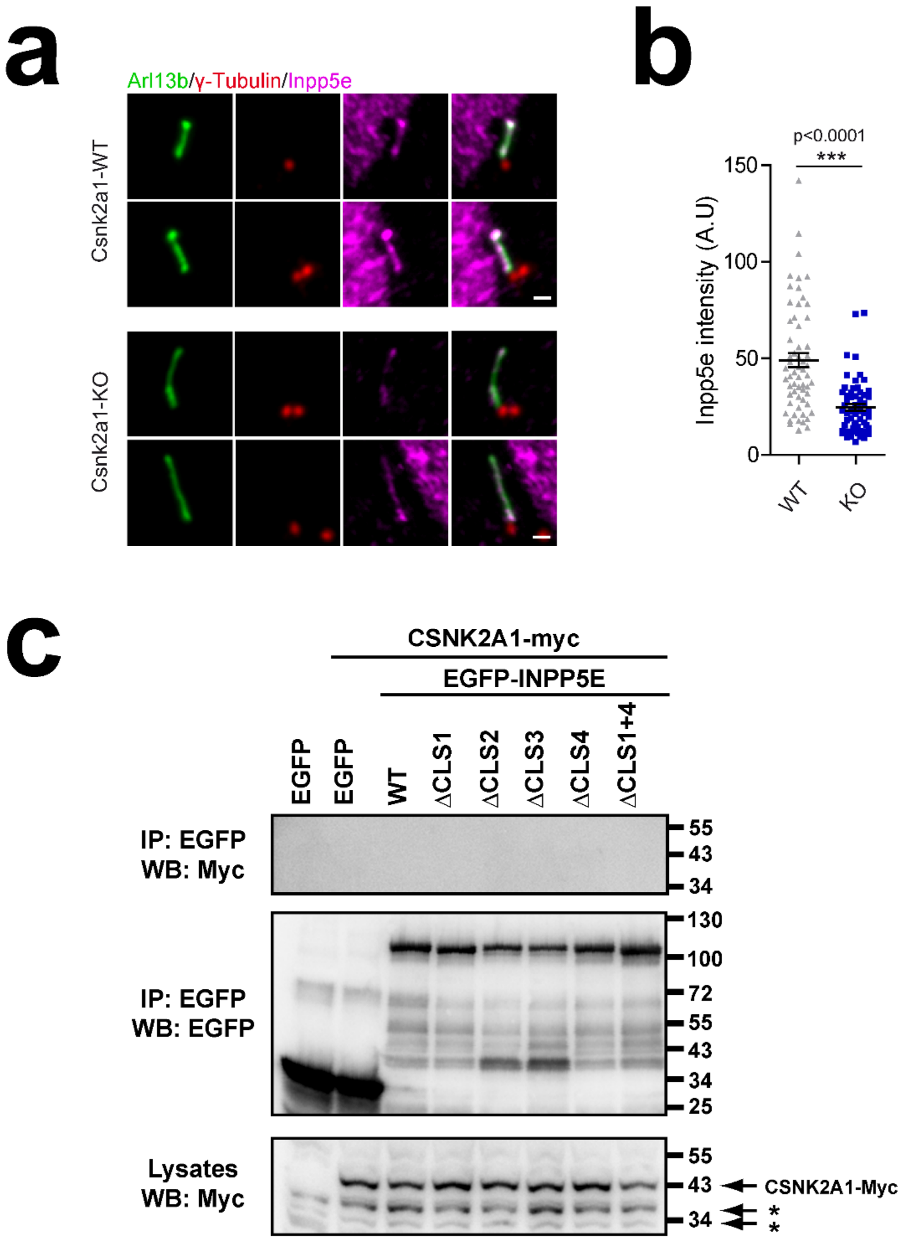


**Figure S6. CSNK2A1 regulates INPP5E ciliary targeting without strongly interacting with it. (a)** Csnk2a1-WT and KO MEFs (Loukil et al. 2021) were stained with antibodies against Arl13b (green), gamma-tubulin (red), and Inpp5e (magenta). Scale bar, 1 µm. **(b)** Quantitation of Inpp5e ciliary intensity in Csnk2a1 WT and KO MEFs. Data are mean ±SEM. Statistical significance in Mann-Whitney non-parametric two-tailed test is shown. **(c)** The indicated EGFP-INPP5E variants were coexpressed in HEK293T cells with CSNK2A1-myc as indicated. Lysates were immunoprecipitated with GFP-Trap beads and analyzed by Western blot with the indicated antibodies. No interaction was detected between EGFP-INPP5E and CSNK2A1-myc. Asterisks indicate non-specific bands. Molecular weight markers are shown on the right.

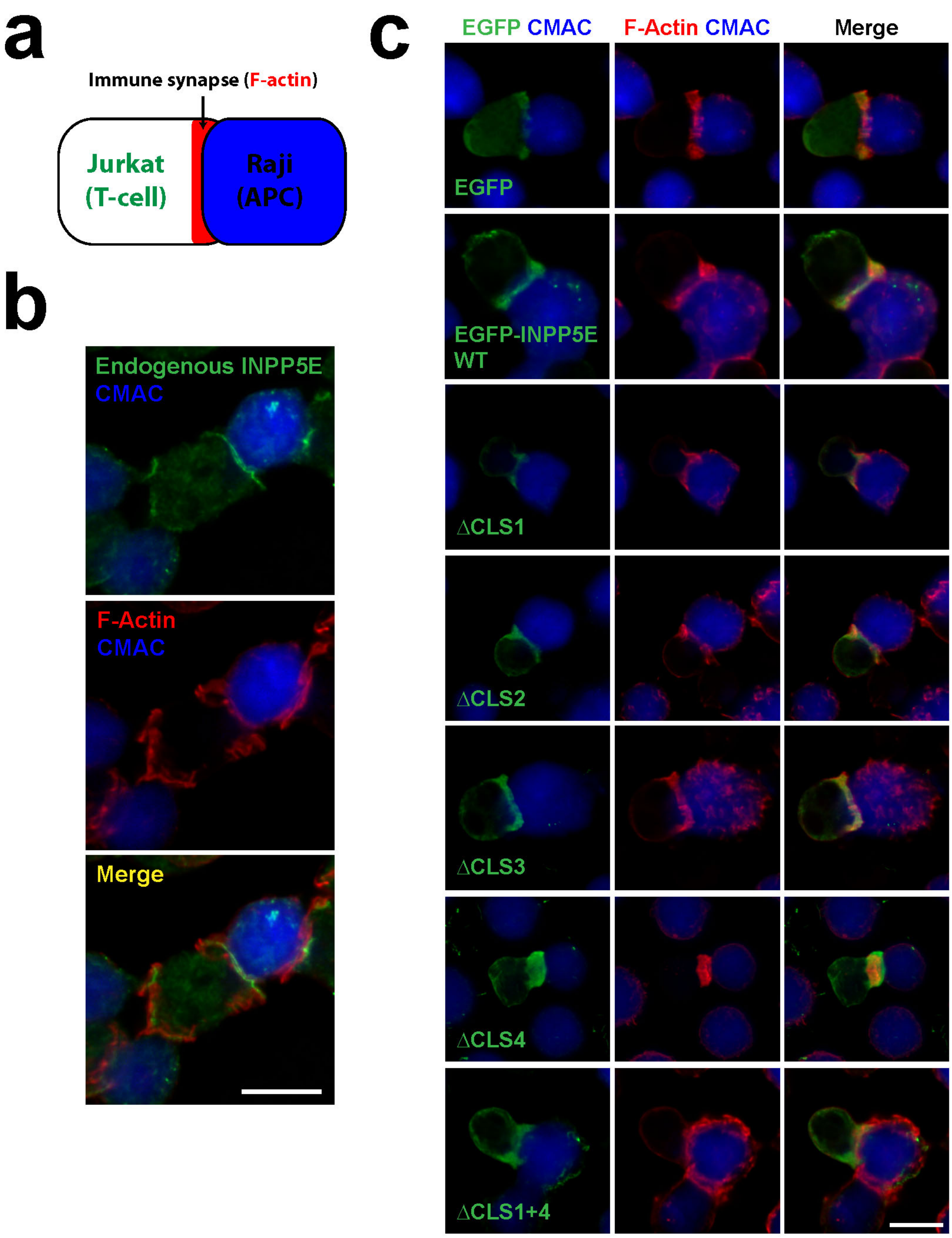


**Figure S7. INPP5E targeting to the T-cell immune synapse is CLS-independent. (a)** Schema depicting an immune synapse (IS) between an antigen-presenting cell (APC) and a T-cell. Herein, Raji and Jurkat cells were used as APC and T-cells, respectively. F-actin is a marker well known to accumulate in productive immune synapses. **(b)** Jurkat cells were challenged with CMAC-labelled SEE-pulsed Raji cells to induce synaptic conjugate formation. Cells were fixed, permeabilized and stained with Alexa Fluor 546-conjugated phalloidin (F-actin, red) and anti-INPP5E antibody (green). CMAC (7-amino-4-

chloromethylcoumarin) is shown in blue. Scale bar, 10 μm. **(c)** Jurkat cells expressing the indicated EGFP fusion proteins were challenged and stained as in (b), except that an anti-EGFP antibody (green) was used instead of anti-INPP5E. Scale bar, 10 μm. Cells in (b-c) were imaged by epifluorescence microscopy.
